# Eos promotes T_H_2 differentiation by propagating the IL-2/STAT5 signaling pathway

**DOI:** 10.1101/2022.11.02.514868

**Authors:** Jasmine A. Tuazon, Kaitlin A. Read, Bharath K. Sreekumar, Michael J. Yaeger, Sanjay Varikuti, Devin M. Jones, Robert T. Warren, Michael D. Powell, Mustafa N. Rasheed, Elizabeth G. Duncan, Lauren M. Childs, Kymberly M. Gowdy, Kenneth J. Oestreich

## Abstract

The Ikaros zinc finger transcription factor Eos has been commonly implicated in regulatory T cells to promote their immunosuppressive functions. Paradoxically, a new role is emerging for Eos in promoting pro-inflammatory responses of conventional CD4^+^ T cells in the dysregulated setting of autoimmunity. Even so, the precise role of Eos in regulating the differentiation and function of healthy effector CD4^+^ T cell subsets remains unclear. Here, we find that Eos is a positive regulator of CD4^+^ T helper 2 (T_H_2) cells—effector T cells implicated in the induction of allergic asthma. Using murine in vitro T_H_2 cells and an in vivo house dust mite asthma model, we found that Eos-deficient T cells had reduced expression of key T_H_2 transcription factors, effector cytokines, and differentiation receptors. Mechanistically, among various T_H_2-polarizing pathways, the IL-2/STAT5 axis and its downstream T_H_2 gene targets emerged as one of the most significantly downregulated networks in Eos deficiency. Using in vitro T_H_2 cells and overexpression of Eos zinc-finger-domain mutants, we discovered that Eos forms a novel complex with and supports the tyrosine-phosphorylated signaling activity of STAT5. Overall, these data define a novel regulatory mechanism whereby Eos promotes IL-2/STAT5 activity to facilitate T_H_2 differentiation.

## Introduction

In response to viruses, bacteria, or parasites in the environment, naïve CD4^+^ T cells differentiate into specialized ‘helper’ subsets that target specific types of pathogens and prime other immune cells’ responses. Because of their pro-inflammatory nature, T helper subset dysregulation can favor the pathogenesis of allergic and autoimmune conditions. T helper 2 (T_H_2) cells are well-known for this dichotomy; while their effector cytokines, such as interleukin-4 (IL-4) and IL-13, are capable of orchestrating the killing or expelling extracellular pathogens (1–3), T_H_2 cells can also induce the pathogenesis of allergic asthma (4, 5). Given this balance between benefit and harm, understanding the regulators of T_H_2 differentiation is of key importance to the prevention and management of T_H_2-mediated diseases.

For CD4^+^ T helper cells to differentiate, naïve CD4^+^ T cells receive three main signals: 1) T cell receptor activation, 2) co-stimulation, and 3) environmental cytokine binding (6, 7). This third signal leads to Janus kinase (JAK)-mediated tyrosine phosphorylation of Signal Transducer and Activator of Transcription (STAT) proteins, causing them to dimerize, translocate to the nucleus, and bind to target T helper subset genes, including lineage-defining transcription factors that commit the cell to a specific T helper subset fate. For a cell to polarize toward the T_H_2 cell fate, IL-4 and STAT6 drive expression of the T_H_2 lineage-defining transcription factor Gata3; however, several studies have also described IL-4/STAT6-independent mechanisms of T_H_2 differentiation (8–11). Further, a second cytokine-STAT pathway, IL-2/STAT5, remains essential for the proliferation, polarization, and effector function of T_H_2 cells (12–14). STAT5 collaborates with Gata3 at critical T_H_2 cytokine gene loci, upregulates CD25 (IL-2Rα) and CD124 (IL-4Rα) receptors to amplify T_H_2 gene program, and stimulates T cell growth (15–20). This IL-2/STAT5 pathway further supports T_H_2 differentiation by directly suppressing alternative T cell programs, including T_FH_ and T_H_17 cells (21–23), and by indirectly upregulating Gata3 to secure the T_H_2 gene program (24–27). Thus, although STAT5 is a well-established player in T_H_2 differentiation, its interplay with wider transcriptional networks in this process is incompletely understood.

A second group of transcription factors that has a close connection with T cell differentiation is the Ikaros zinc finger (IkZF) family. Its five members include Ikaros (*Ikzf1*), Helios (*Ikzf2*), Aiolos (*Ikzf3*), Eos (*Ikzf4*), and Pegasus (*Ikzf5*). These regulators have conserved N-terminal DNA-binding domains and C-terminal protein-interaction domains that enable IkZFs to mediate T and B lymphocyte hematopoiesis (28); IkZF factors recruit chromatin remodeling complexes that can either promote or repress T cell development and differentiation programs (29). For example, Ikaros has been shown to promote differentiation of T_H_2 (30, 31), T_H_17 (32–34), T_FH_ (35, 36), and T_REG_ subsets (31–33, 37, 38). Helios stabilizes a Foxp3+ T_REG_ immunosuppressive phenotype and is associated with—but not required for—T_H_2 and T_FH_ differentiation (39–42). Meanwhile, Aiolos has been shown to promote T_H_17 and T_FH_ cell fates by repressing IL-2 (21, 43–45). Furthermore, our lab has previously shown that Aiolos cooperatively binds with STAT3 to upregulate Bcl-6 expression (36). However, whether such an IkZF-STAT relationship exists among other family members, as well as the mechanisms by which IkZFs interact with and modulate STAT proteins to regulate T cell differentiation, remains incompletely understood.

To this end, our laboratory investigated these questions in the context of the fourth IkZF family member, Eos, which holds a complex role in the differentiation of T cell subsets that are promoted by opposing STAT pathways. Eos has been largely described in T_REG_ differentiation, wherein it complexes with Foxp3 to support T_REG_ differentiation, immunosuppressive function, and IL-2 repression (46–49). Despite this, some work has cast doubt on the strict requirement for Eos in the development of robust T_REG_ suppressor function (50), indicating that more investigation is needed on the exact mechanisms of Eos’ regulation in T cell polarization. Indeed, Eos’ role in inflammatory populations remains ill-defined, as Eos seems to activate some CD4^+^Foxp3^-^ T_CONV_ cells (46, 50) while repressing other cells such as T_H_17 cells (51). Given Eos’ enigmatic role in differentiation, our lab sought to define the mechanistic underpinnings that can explain Eos’ dual contributions to immunosuppressive and pro-inflammatory T cells.

In the present work, we show that Eos is a novel positive regulator of pro-inflammatory T_H_2 cells. Our in vitro-generated T_H_2 cells and in vivo murine asthma models demonstrate that in Eos deficiency, gene and protein expression of key T_H_2 transcription factors, cytokine receptors, and effector cytokines are significantly reduced. These experiments further indicated that Eos and IL-2/STAT5, an essential pathway to T_H_2 differentiation, amplify each other’s expression in a positive-feedback loop. Strikingly, we found that, in both T_H_2 and T_REG_ cells, Eos forms a direct complex with and increases the activity of STAT5 to regulate expression of downstream T_H_2 differentiation genes, such as *Il4ra*. Taken together, our data reveal a novel mechanism by which Eos binds to and positively regulates STAT5 tyrosine-phosphorylated signaling activity to promote T_H_2 differentiation and effector cytokine production.

## Materials and Methods

### Mouse strain and cell lines

C57BL/6J mice were purchased from the Jackson Laboratory. *Ikzf4^-/-^* mice were generously provided by Drs. Ethan Shevach (NIH) and Bruce Morgan (Harvard Medical School) (50). To limit biological variability in in vitro mice studies, male and female mice with age- and sex-matched controls were used to avoid unintentional bias. Due to known asthma pathogenesis variability associated with estrogen (52–54), we used male mice for the in vivo murine house dust mite asthma model. The Institutional Animal Care and Use Committees of The Ohio State University approved all experiments involving the use of mice. All methods were performed in accordance with the approved guidelines.

EL4 cells were acquired from the American Type Culture Collection (TIB-39) and cultured in complete RPMI (RPMI-1640, 10% FBS, 1% Pen/Strep).

### CD4^+^ T cell isolation and culture

Naive CD4^+^ T cells were isolated from the spleens and lymph nodes of 5-8 week old mice using the BioLegend MojoSort naïve CD4^+^ T cell isolation kit according to the manufacturer’s recommendations. Naïve cell purity was verified by flow cytometry and routinely exceeded 96-98%. Harvested naïve CD4^+^ T cells were plated at a density of 1.5 × 10^5^ cells/mL in 2 ml of complete IMDM ((IMDM [Life Technologies], 10% FBS [26140079, Life Technologies], 1% Penicillin-Streptomycin [Life Technologies], and 0.05% (50μM) 2-ME [Sigma-Aldrich]). Cells were stimulated on plate-bound anti-CD3 (clone 145-2C11; 5μg/mL) and anti-CD28 (clone 37.51; 2μg/mL) in the presence T_REG_, T_H_2, T_FH_, or T_H_17 polarizing conditions (**Table 1**). Blocking antibodies were added on day 0 at plating, and polarizing cytokines were added 24 hours later. Cells were split on day 3 and plated at 2.5 × 10^5^ cells/ml in 2 ml, replated with polarizing cytokines, and left to incubate before harvesting at day 5.

**Table 1.**
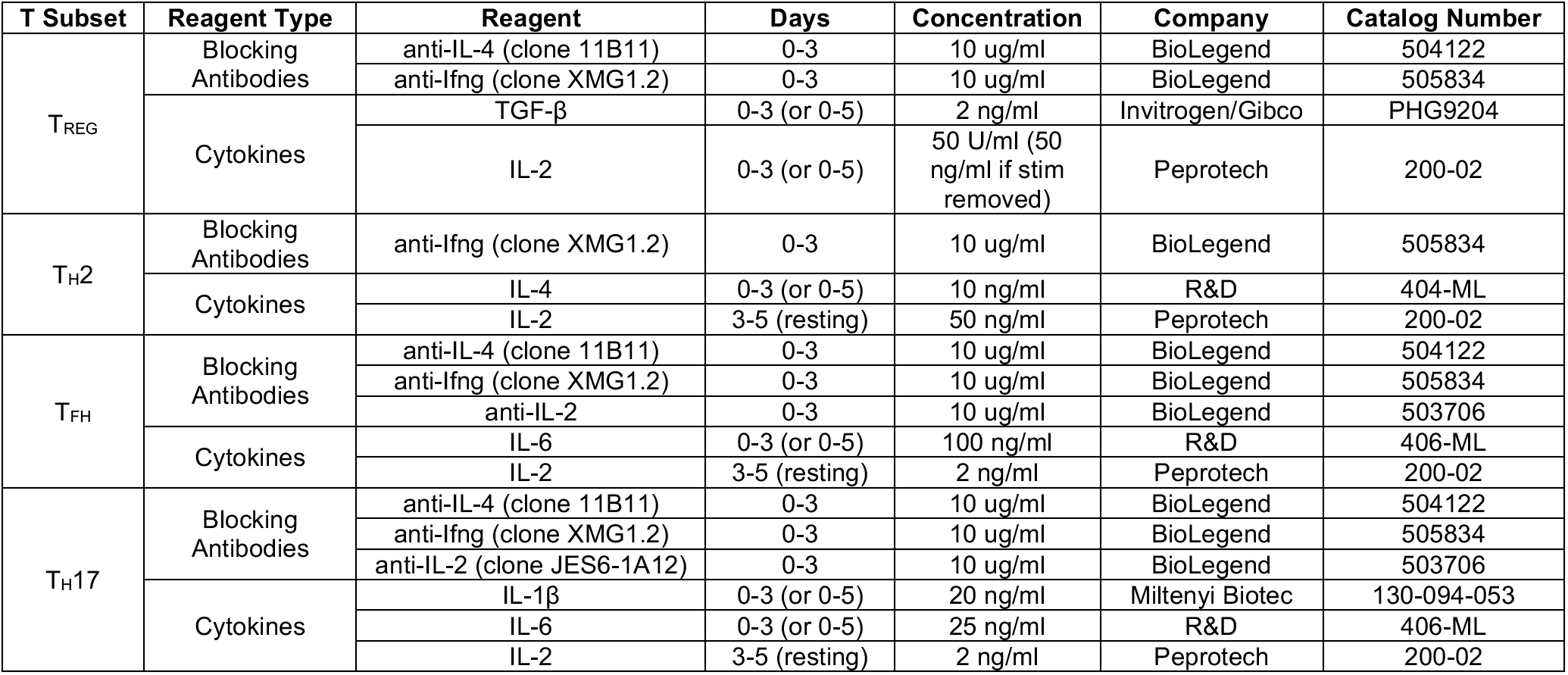
In vitro T subset polarizing conditions

| T Subset | Reagent Type | Reagent | Days | Concentration | Company | Catalog Number |
| --- | --- | --- | --- | --- | --- | --- |
| T<sub>REG</sub> | Blocking Antibodies | anti-IL-4 (clone 11B11) | 0-3 | 10 ug/ml | BioLegend | 504122 |
|  |  | anti-Ifng (clone XMG1.2) | 0-3 | 10 ug/ml | BioLegend | 505834 |
|  | Cytokines | TGF-β | 0-3 (or 0-5) | 2 ng/ml | Invitrogen/Gibco | PHG9204 |
|  |  | IL-2 | 0-3 (or 0-5) | 50 U/ml (50 ng/ml if stim removed) | Peprtech | 200-02 |
| T<sub>H2</sub> | Blocking Antibodies | anti-Ifng (clone XMG1.2) | 0-3 | 10 ug/ml | BioLegend | 505834 |
|  | Cytokines | IL-4 | 0-3 (or 0-5) | 10 ng/ml | R&D | 404-ML |
|  |  | IL-2 | 3-5 (resting) | 50 ng/ml | Peprtech | 200-02 |
| T<sub>FH</sub> | Blocking Antibodies | anti-IL-4 (clone 11B11) | 0-3 | 10 ug/ml | BioLegend | 504122 |
|  |  | anti-Ifng (clone XMG1.2) | 0-3 | 10 ug/ml | BioLegend | 505834 |
|  |  | anti-IL-2 | 0-3 | 10 ug/ml | BioLegend | 503706 |
|  | Cytokines | IL-6 | 0-3 (or 0-5) | 100 ng/ml | R&D | 406-ML |
|  |  | IL-2 | 3-5 (resting) | 2 ng/ml | Peprtech | 200-02 |
| T<sub>H17</sub> | Blocking Antibodies | anti-IL-4 (clone 11B11) | 0-3 | 10 ug/ml | BioLegend | 504122 |
|  |  | anti-Ifng (clone XMG1.2) | 0-3 | 10 ug/ml | BioLegend | 505834 |
|  |  | anti-IL-2 (clone JES6-1A12) | 0-3 | 10 ug/ml | BioLegend | 503706 |
|  | Cytokines | IL-1β | 0-3 (or 0-5) | 20 ng/ml | Miltenyi Biotec | 130-094-053 |
|  |  | IL-6 | 0-3 (or 0-5) | 25 ng/ml | R&D | 406-ML |
|  |  | IL-2 | 3-5 (resting) | 2 ng/ml | Peprtech | 200-02 |

### Overexpression experiments

Nucleofection assays in EL4 T cells were performed with the Lonza 4D Nucleofection system (buffer SF, program CM-120). Expression vectors were made by cloning the indicated coding sequences into the pcDNA3.1/V5-His-TOPO vector (cat. no. K480001; Life Technologies). Constitutively active STAT5B (STAT5B^CA^) was constructed as previously described (55). In brief, point mutations were generated using the Agilent QuikChange Site-Directed Mutagenesis Kit (part number 200519) according to the manufacturer’s instructions. Wildtype and mutant coding sequences were transferred to the pEF1/V5-His vector (cat. no. V920; Life Technologies) for overexpression. EL4s were transfected for 22-24 hours before being harvested for downstream analysis. Overexpression of proteins was assessed via immunoblot using both V5 tag- and protein-specific antibodies, and alterations in gene expression were assessed via qRT-PCR analysis.

### Promoter-reporter analysis

An *Ikzf4* promoter-reporter construct (pGL3-Ikzf4) was generated by cloning the regulatory region of *Ikzf4* (2 kb upstream of the transcriptional start site) into the pGL3-Basic vector (Promega). EL4 T cells were nucleofected in combination with either a STAT5B^CA^, STAT5B^WT^, or STAT3^CA^ expression vectors. SV40-*Renilla* vector control was used to assess transfection efficiency. Following 22-24 hours of recovery, samples were harvested and luciferase expression was analyzed using the Dual-Luciferase Reporter Assay System according to the manufacturer’s instructions (Promega).

### RNA isolation and qRT-PCR

Total RNA was isolated from the indicated cell populations using the Macherey-Nagel Nucleospin RNA Isolation kit as recommended by the manufacturer. cDNA was generated from mRNA template using the Superscript IV First Strand Synthesis System with provided oligo dT primer (Thermo Fisher). qRT-PCR reactions were performed with the SYBR Select Mastermix for CFX (ThermoFisher) using 6 ng cDNA per reaction and the primers provided below (**Table 2**). All qRT-PCR was performed on the CFX Connect (BioRad). Data were normalized to *Rps18* and presented either relative to *Rps18* or relative to the control sample, as indicated.

**Table 2.**
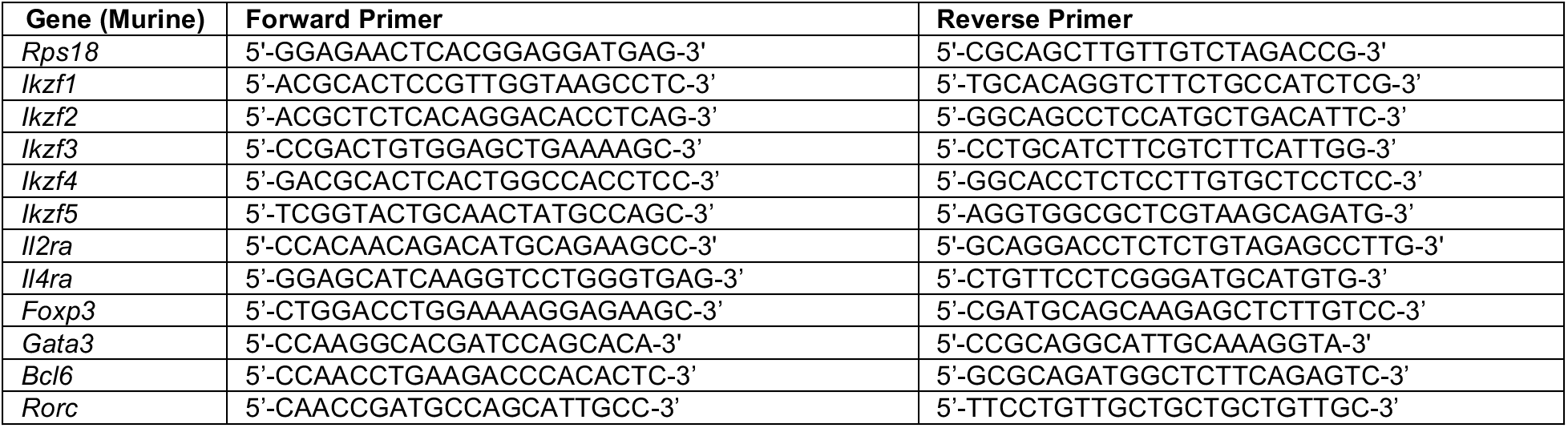
qRT-PCR Primers

**Table 3.**
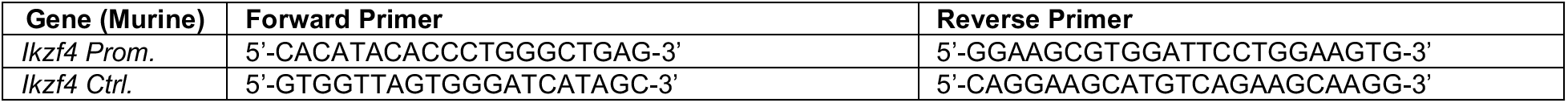
ChIP Primers

### Immunoblot analysis

An equivalent number of cells were harvested for each analysis, lysed directly in 1X SDS loading dye (50 mM Tris [pH 6.8], 100 mM DTT, 2% SDS, 0.1% bromophenol blue, 10% glycerol) and boiled for 15 minutes. Lysates were separated via SDS-PAGE and transferred to 0.45μm nitrocellulose membrane. Membranes were blocked with 2% nonfat dry milk in 1X TBST (10 mM Tris [pH 8], 150 mM NaCl, 0.05% Tween-20), and detection of indicated proteins was carried out using the following antibodies: Eos (W15169A; BioLegend, 1:500), pY-STAT5 (BD Biosciences, 611964, 1:5000), STAT5 (Cell Signaling, 94205S, 1:5000), STAT5B (1:5000; Santa Cruz 1656), β-actin (GenScript, 1:15,000), goat anti-mouse (Jackson Immunoresearch, 1:5000-1:10,000), mouse anti-rabbit (Santa Cruz, 1:15,000).

### Co-immunoprecipitation (Co-IP)

Co-immunoprecipitation analyses were performed using primary murine in vitro-generated T_H_2 cells, or EL4 cells overexpressing indicated proteins as previously described (36, 56, 57). Briefly, lysates were immunoprecipitated with anti-STAT5 (0.785μg /IP, 94205S, Cell Signaling) or an isotype control antibody overnight at 4°C. The following day, samples were incubated in the presence of Protein A Sepharose beads (Millipore) for 1-2h, and immunoprecipitated proteins were analyzed via subsequent immunoblot analysis. Antibodies to detect immunoprecipitated proteins were as follows: Eos (1:500, Biolegend, W15169A), STAT5 (1:5000; Santa Cruz, 74442x), and V5 (1:20,000; R960-25, Invitrogen).

### RNA-seq analysis

Naïve CD4^+^ T cells were cultured under T_H_2-polarizing conditions for 3 days. Total RNA was isolated using the Macherey-Nagel Nucleospin kit according to the manufacturer’s instructions. Samples were provided to Azenta Life Sciences for polyA selection, library preparation, sequencing, and DESeq2 analysis. Genes with a p-value < 0.05 were considered significant, and those with an absolute log2 fold change of ≥1.0 were defined as differentially expressed genes (DEGs) for each comparison in the present study. Genes pre-ranked by multiplying the sign of the fold-change by −log10 (p-value) were analyzed using the Broad Institute Gene Set Enrichment Analysis (GSEA) software for comparison against ‘hallmark’, ‘gene ontology’, and ‘immunological signature’ gene sets. Heatmap generation and clustering (by Euclidean distance) were performed using normalized log2 counts from DEseq2 analysis and the Morpheus software (https://software.broadinstitute.org/morpheus). Volcano plots were generated using −log10(p-value) and log2 fold change values from DEseq2 analysis and VolcaNoseR software (https://huygens.science.uva.nl/VolcaNoseR/) (58).

### HDM sensitization and challenge

On days 0 and 7, we sensitized C57BL/6J mice with 10 μg of HDM solution (*Dermatophagoides pteronyssinus*, 28,750 endotoxin units/vial; catalog number XPB82D3A2.5, Greer Laboratories Inc.) in 50 μl of PBS via oropharyngeal (o.p.) aspiration. Then, on days 14, 15, and 16, we challenged the mice o.p. with 2 μg of the HDM preparation. On day 17, lungs were harvested for downstream T cell analysis.

### Flow cytometry

For panels requiring analysis of cytokine production, cells were first incubated in complete IMDM with PMA/ionomycin and protein transport inhibitors (PTI) for 4 hours. For extracellular staining, samples were pre-incubated for 5 minutes at 4°C with TruStain FcX™ (anti-mouse CD16/32) Fc block (clone 93; BioLegend). Samples were then stained for extracellular markers in the presence of Fc block for 30 minutes at 4°C protected from light using the following antibodies: CD3 (APC/Cy7; 1:300; clone 17A2; BioLegend), CD4 (AF488; 1:300 (for cells not given PMA/ionomycin and PTI) or 1:100 (for cells incubated with PMA/ionomycin and PTI); clone GK1.5; R&D Systems); CD44 (V450; 1:300; clone IM7; BD Biosciences); CD62L (APC-eFluor 780; 1:300; clone MEL-14; ThermoFisher); CD25 (APC; 1:300; clone PC61.5; ThermoFisher); and Ghost Dye (V510; 1:400; Tonbo Biosciences). Cells were then washed twice with FACS buffer prior to intracellular staining. For intracellular staining, cells were fixed and permeabilized using the eBioscience Foxp3 transcription factor staining kit (ThermoFisher) for 30 minutes at room temperature, or overnight at 4°C. Following fixation, samples were stained with the following antibodies in 1X eBioscience permeabilization buffer for 1 hour at room temperature protected from light: Foxp3 (1:100; clone FJK-16s; ThermoFisher); Gata3 (PE-Cy7; 1:20; clone TWAJ; ThermoFisher); IL-4 (APC; 1:50; clone 11B11; BioLegend); IL-13 (PE; 1:50; clone W17010B; BioLegend); Cells were washed with 2X eBioscience permeabilization buffer and resuspended in FACS buffer for analysis. Where appropriate, fluorescence-minus-one (FMO) controls were used to determine nonspecific staining. Samples were run on a BD FACS Canto II flow cytometer and analyzed using FlowJo software (version 10.8.1).

### Chromatin immunoprecipitation (ChIP)

ChIP assays were performed as described previously (59). Resulting chromatin fragments were immunoprecipitated with antibodies against STAT5 (R&D AF2168, 10 μg/IP) or IgG control (Abcam ab6709; 10 μg/IP, matched to experimental antibody). Enrichment of the indicated proteins was analyzed via qPCR with the primers below. Samples were normalized to total DNA controls and isotype control antibodies were used to ensure lack of non-specific background.

### Statistical analysis

All statistical analyses were performed using the GraphPad Prism software (version 9.4.1). For single comparisons, unpaired Student’s *t*-tests were performed. For multiple comparisons, one-way ANOVA with Tukey’s multiple comparison tests were performed. Error bars indicate the standard error of the mean. *P* values <0.05 were considered statistically significant.

## Results

### 1. Eos expression is elevated in T_H_2 cells and is promoted by IL-2/STAT5 signaling

Our lab has previously shown that the IkZF Aiolos is elevated in T_FH_ cells and aides in their differentiation by cooperating with STAT3 to upregulate Bcl-6 expression (36). To screen for IkZFs upregulated in other CD4^+^ T helper subsets, we isolated naïve wild-type (WT) CD4^+^ T cells from C57BL/6J mice and cultured them under various T subset polarizing conditions, including T_H_2- and T_FH_-polarizing conditions. Here, we found that, consistent with the established literature (30, 36, 60), Ikaros was the most highly expressed IkZF in T_H_2 and T_FH_ cells (**Supplemental Fig. 1A**). Comparing the four remaining IkZF family members, we found that transcript levels of Eos were elevated in T_H_2 cells compared to other IkZFs (**Fig. 1A**). Additionally, we saw an inverse pattern of Eos and Aiolos gene expression, with *Ikzf3* expression reduced in T_H_2 cells relative to T_FH_ cells and *Ikzf4* transcript elevated in T_H_2 cells compared to T_FH_ cells (**Fig. 1A**). This elevation in Eos in pro-inflammatory T_H_2 cells stood out to us, as majority of studies to date have associated Eos with the immunosuppressive T_REG_ gene program (46–49). As such, we compared Eos gene and protein expression between a range of CD4^+^ T helper cells, including T_REG_, T_H_2, T_FH_, and T_H_17 cells (**Supplemental Fig. 1B,C**). We found that although Eos gene expression was markedly elevated in T_REG_ cells, Eos was still significantly elevated in T_H_2 cells compared to T_FH_ and T_H_17 cells (**Fig. 1B**). We considered the commonalities that exist between T_REG_ and T_H_2 cells—and not T_FH_ or T_H_17 cells—and noted that T_REG_ and T_H_2 cells are promoted by the IL-2/STAT5 axis, while the T_FH_ and T_H_17 gene programs are repressed by this pathway (61).

**Figure 1:**
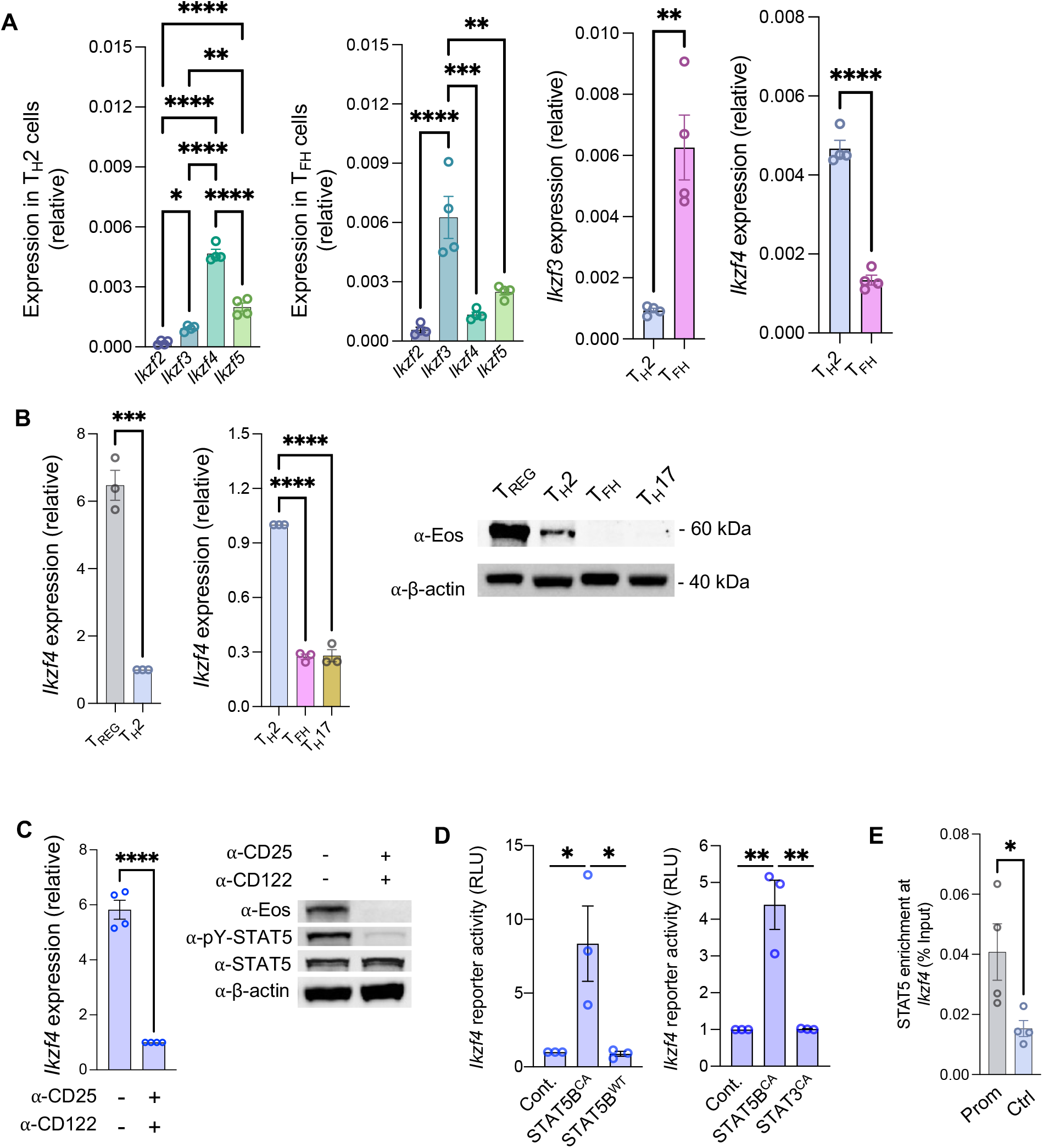
Eos is elevated in T_H_2 cells, and the IL-2/STAT5 pathway regulates Eos expression. (**A**) Naïve WT CD4^+^ T cells from C57BL/6J mice were cultured on plates coated with α-CD3/α-CD28 stimulation under T_H_2- and T_FH_-polarizing conditions (**Table 1**) for 3 days before being removed from stimulation and cultured in resting T_H_2 (IL-4 and IL-2) and T_FH_ (IL-6, IL-2) conditions for an additional 48 hours before harvesting on day 5. RNA was isolated and qRT-PCR was used to assess gene expression. Data were normalized to *Rps18* (n = 4 independent experiments, mean ± s.e.m., \**P* < 0.05, \*\**P* < 0.01, \*\*\**P* < 0.001. \*\*\*\**P* < 0.0001; one-way ANOVA with Tukey’s post-hoc test). **(B)** Cells were cultured as in ‘A’ under T_REG_-, T_H_2-, T_FH_-, and T_H_17-polarizing conditions. RNA was isolated and qRT-PCR was used to assess gene expression. Data were normalized to *Rps18*. An immunoblot was also performed to assess the relative abundance of Eos (n = 4 independent experiments, mean ± s.e.m., \*\*\**P* < 0.001. *\*\*\*\*P* < 0.0001; one-way ANOVA with Tukey’s post-hoc test). **(C)** Cells were cultured as in ‘A’ under T_H_2-polarizing conditions only or with neutralizing antibodies for CD25 (α-IL-2Rα) and CD122 (α-IL-2Rβ). After 4 days, cells were harvested, RNA was isolated, and qRT-PCR was used to assess *Ikzf4* (Eos) expression. Data were normalized to *Rps18* and presented as fold change relative to anti-CD25/anti-CD122 sample. Immunoblot was performed to assess the relative abundance of Eos, pY-STAT5, and STAT5 expression (n = 4 independent experiments, mean ± s.e.m., *\*\*\*\*P* < 0.0001; unpaired Student’s *t*-test). **(D)** EL4 T cells were transfected with an *Ikzf4* promoter-reporter construct in combination with either an empty vector control, constitutively active STAT5B (STAT5B^CA^), STAT5B^WT^ or STAT3^CA^ expression vectors. After 24 hours, cells were harvested for promoter-reporter assay. Luciferase promoter-reporter values were normalized to a *Renilla* control and presented relative to the empty vector control sample (n = 3 independent experiments, mean ± s.e.m., \**P* < 0.05, \*\**P* < 0.01; one-way ANOVA with Tukey’s post-hoc test). **(E)** Chromatin Immunoprecipitation (ChIP) assays to assess STAT5 association with the *Ikzf4* locus in in vitro T_H_2 cells that were cultured, split at day 3, and harvested at day 5 as in ‘A’. The *Ikzf4* promoter (“Prom”) and an upstream control region (“Ctrl”) were interrogated for STAT5 enrichment. Data are presented as percent enrichment relative to a “total” input sample. ChIP analysis was performed using anti-STAT5 and anti-IgG control antibodies for the indicated regions. Data are normalized to total input sample. IgG values were subtracted from percent enrichment (*n* = 4 independent experiments, mean ± s.e.m., * *P* < 0.05; unpaired Student’s *t*-test).

To then assess how the IL-2/STAT5 pathway could be affecting Eos expression, we first disrupted IL-2 signaling by culturing T_H_2 cells with both anti-CD25 (anti-IL-2Rα) and anti-CD122 (anti-IL-2Rβ) blocking antibodies (**Fig. 1C**). We found that blocking IL-2 signaling significantly attenuated Eos transcript and protein expression (**Fig. 1C**), indicating that signals downstream of IL-2 positively regulate Eos expression in T_H_2 cells. We then designed a constitutively active STAT5B (STAT5B^CA^) vector—choosing STAT5B over STAT5A because STAT5B plays a more dominant role in immune cells (62)—and used a promoter-reporter construct containing the Eos (*Ikzf4*) promoter in the presence and absence of STAT5B^CA^ (**Fig. 5B**). Indeed, we observed a significant increase in *Ikzf4* promoter activity following overexpression of STAT5B^CA^ (**Fig. 1D**). Supporting the specificity of these findings to STAT5B^CA^, we did not observe an increase in *Ikzf4* promoter activity in response to overexpression of STAT3^CA^ or wild-type STAT5B (STAT5B^WT^) (**Fig. 1D**). Finally, to determine whether STAT5 binds to the endogenous *Ikzf4* promoter, we performed chromatin immunoprecipitation (ChIP) analysis and found that STAT5 enrichment was significantly increased at the *Ikzf4* promoter in T_H_2 cells (**Fig. 1E**). Collectively, these findings show that IL-2/STAT5 signaling promotes Eos expression in T_H_2 cells.

### 2. The T_H_2-transcriptional program is disrupted in Eos deficiency

Given this elevated Eos expression specific to T_H_2 cells, we then asked whether Eos promotes T_H_2 gene patterns underlying differentiation and effector cytokine production. To do this, we cultured WT and Eos-deficient naïve CD4^+^ T cells under T_H_2-polarizing conditions for 3 days and performed bulk RNA-seq analysis (**Fig. 2A**). Heatmap analyses revealed global transcriptomic changes with Eos deficiency (**Fig. 2B**), with RNA-seq and confirmatory qRT-PCR analyses showing significant downregulation of T_H_2 differentiation genes, including key T_H_2 transcription factors *Gata3* and *Prdm1* (which encodes Blimp-1, a factor that protects the T_H_2 gene program), effector cytokines *Il4, Il5*, and *Il13*, and cell surface receptor *Il4ra* (**Fig. 2C-D, Supplemental Fig. 2A**). Conversely, lineage-defining transcription factors *Bcl6* and *Rorc* and other markers associated with the T_FH_ and T_H_17 gene programs were upregulated (**Fig. 2C-D**). Gene Set Enrichment Analysis (GSEA) of curated immunologic signature gene sets in these in vitro-generated T_H_2 cells revealed that gene sets typically upregulated in IL-4-treated naïve CD4^+^ T cells were downregulated in Eos-deficient T_H_2 cells (**Fig. 2E**). To contrast, gene sets associated with T_H_17- and T_FH_-polarizing conditions were upregulated (**Supplemental Fig. 2B**). Taken together, these data indicate that the T_H_2 transcriptional program is compromised in Eos-deficient CD4^+^ T cells.

**Figure 2.**
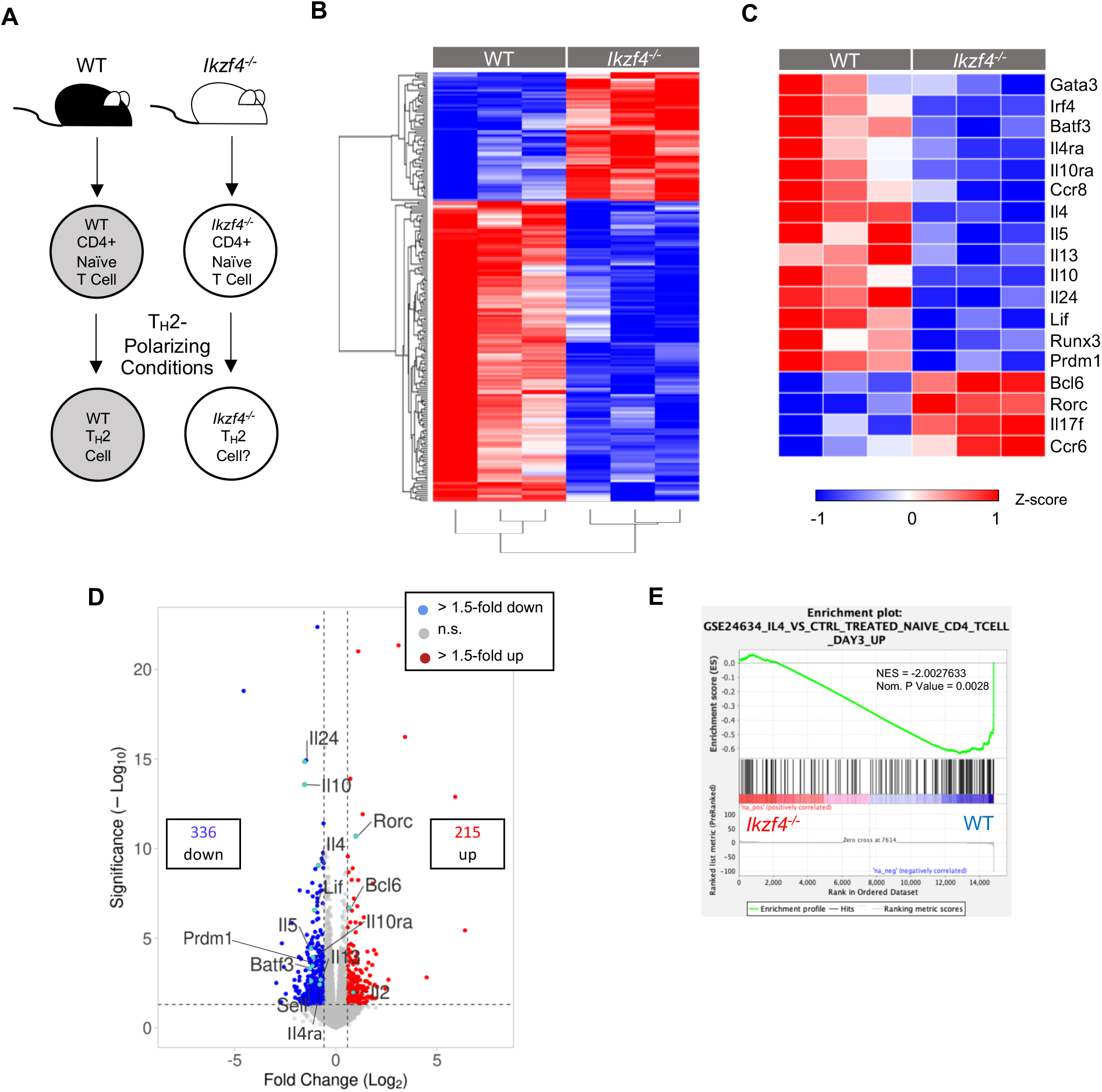
Eos deficiency results in global T_H_2 transcriptional changes. **(A)** Naïve CD4^+^ T cells from WT and Eos-deficient C57BL/6J mice were cultured on plates coated with α-CD3/α-CD28 stimulation under T_H_2-polarizing conditions (**Table 1**) and harvested on day 3. (**B**) RNA-seq analysis was performed to assess differentially expressed genes (DEGs) between WT and Eos-deficient cells. (**C**) Heatmap of DEGs positively and negatively associated with the T_H_2 gene program in WT vs. Eos-deficient cells. Changes in gene expression are presented as relative expression by row (gene). **(D)** Volcano plot displaying gene expression changes in Eos-deficient vs. WT cells. Genes of particular interest are labeled. Genes were color-coded as follows: no significant changes in expression (gray), upregulated genes with > 1.5-fold change in expression with a *P* < 0.05 (red), downregulated genes with > 1.5-fold change in expression with a *P* < 0.05 (dark blue), selected genes of interest accompanying the heatmap (teal blue). **(E)** Pre-ranked (sign of fold change x −log_10_(p-value)) genes were analyzed using the Broad Institute Gene Set Enrichment Analysis (GSEA) software for comparison against ‘immunological signature’ gene sets. Data are compiled from 3 biological replicates from 3 independent experiments.

### 3. The IL-2/STAT5 pathway is downregulated in Eos-deficient T_H_2 cells

Our transcriptomic data also revealed a downregulation of genes associated with the IL-2/STAT5 pathway in Eos-deficient T_H_2 cells. Specifically, RNA-seq showed downregulation of *Il2ra, Il2rb, Jak3*, and *Stat5b* (**Fig. 3A**). To contrast, IL-2 was upregulated, suggesting that the downregulation of the downstream components of the pathway were not due to a loss in IL-2 (**Fig. 3A**). GSEA further revealed downregulation of the hallmark IL-2/STAT5 signaling gene set (**Fig. 3B**). To our surprise, we noted that gene sets associated with STAT tyrosine phosphorylation at Y694/99—modifications most closely associated with STAT dimerization, nuclear translocation, and transcriptional activity (63, 64)—were also downregulated with the loss of Eos (**Fig. 3C**). Accordingly, immunoblot of WT and Eos-deficient T_H_2 cells revealed a reduction in the levels of tyrosine-phosphorylated STAT5 (pY-STAT5) (**Fig. 3D**). Notably, this change in pY-STAT5 was also observed in in vitro-generated T_REG_ cells (**Supplemental Fig. 2C**), suggesting this relationship between Eos and IL-2/STAT5 may be conserved across cell types. Together, these data demonstrate that Eos deficiency is associated with downregulated IL-2 receptor expression and reduced STAT5 activity.

**Figure 3.**
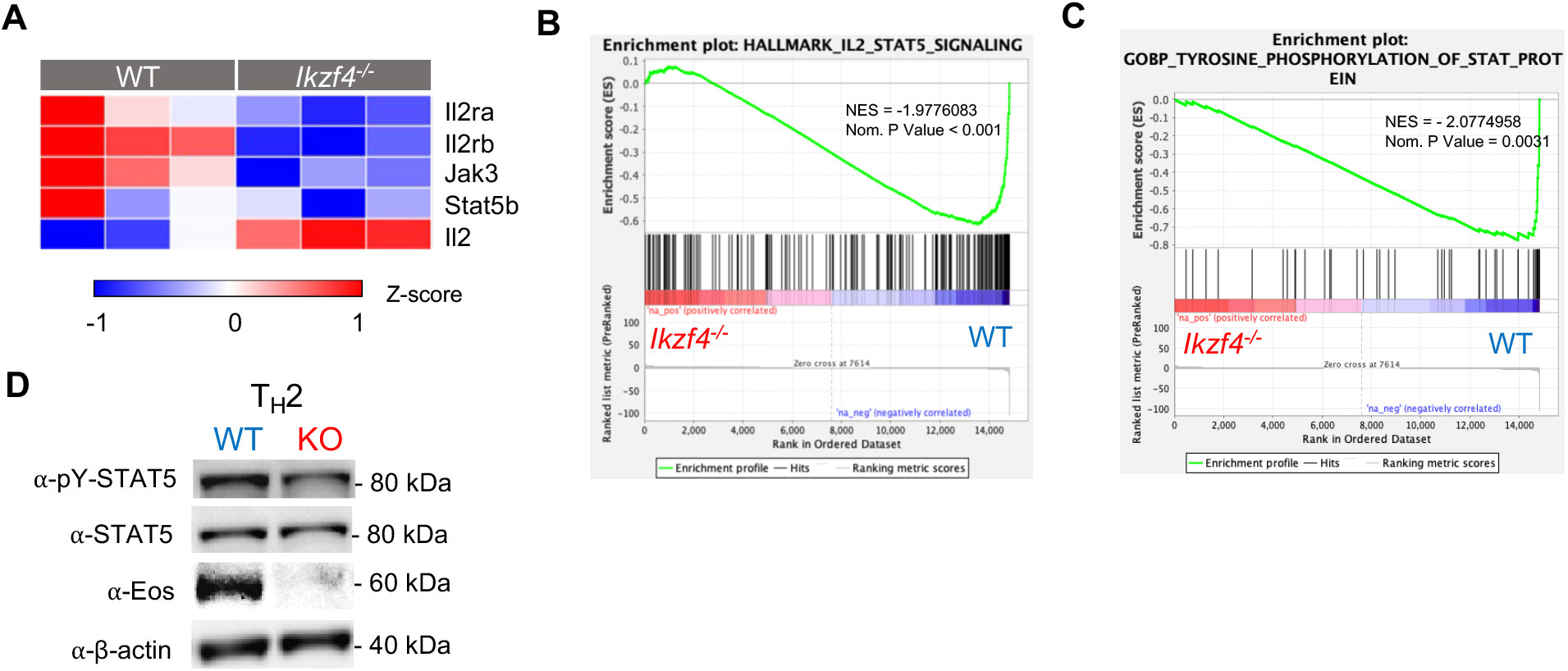
The IL-2/STAT5 pathway is positively regulated by Eos in in vitro T_H_2 cells. **(A)** Heatmap of IL-2/STAT5 pathway DEGs both positively and negatively associated with the T_H_2 gene program in WT vs. Eos-deficient cells; changes in gene expression are presented as relative expression by row (gene). **(B,C)** Pre-ranked (sign of fold change x −log_10_(p-value)) genes were analyzed using the Broad Institute GSEA software for comparison against ‘hallmark’ and ‘gene ontology’ gene sets. Data are compiled from 3 biological replicates from 3 independent experiments. **(D)** Immunoblot of day 3 WT vs. Eos-deficient T_H_2 cells was performed to assess the relative abundance of pY-STAT5, STAT5, and Eos expression (n = 3 independent experiments).

### 4. Eos-deficient mice have dysregulated T_H_2 differentiation and effector cytokine production

We next aimed to analyze how T_H_2 differentiation and effector cytokine production may be impacted in the context of an immune response. We decided to implement a murine house dust mite (HDM) allergic asthma model—a robust and well-established model of T_H_2-induced asthma (65). On days 0 and 7, we sensitized C57BL/6J mice with 10 μg of HDM in sterile PBS via oropharyngeal (o.p.) aspiration (**Fig. 4A**). Then, on days 14, 15, and 16, we challenged the mice o.p. with 2 μg of the HDM preparation. Finally, we harvested lungs on day 17 and isolated lymphocytes for flow cytometric analysis.

**Figure 4.**
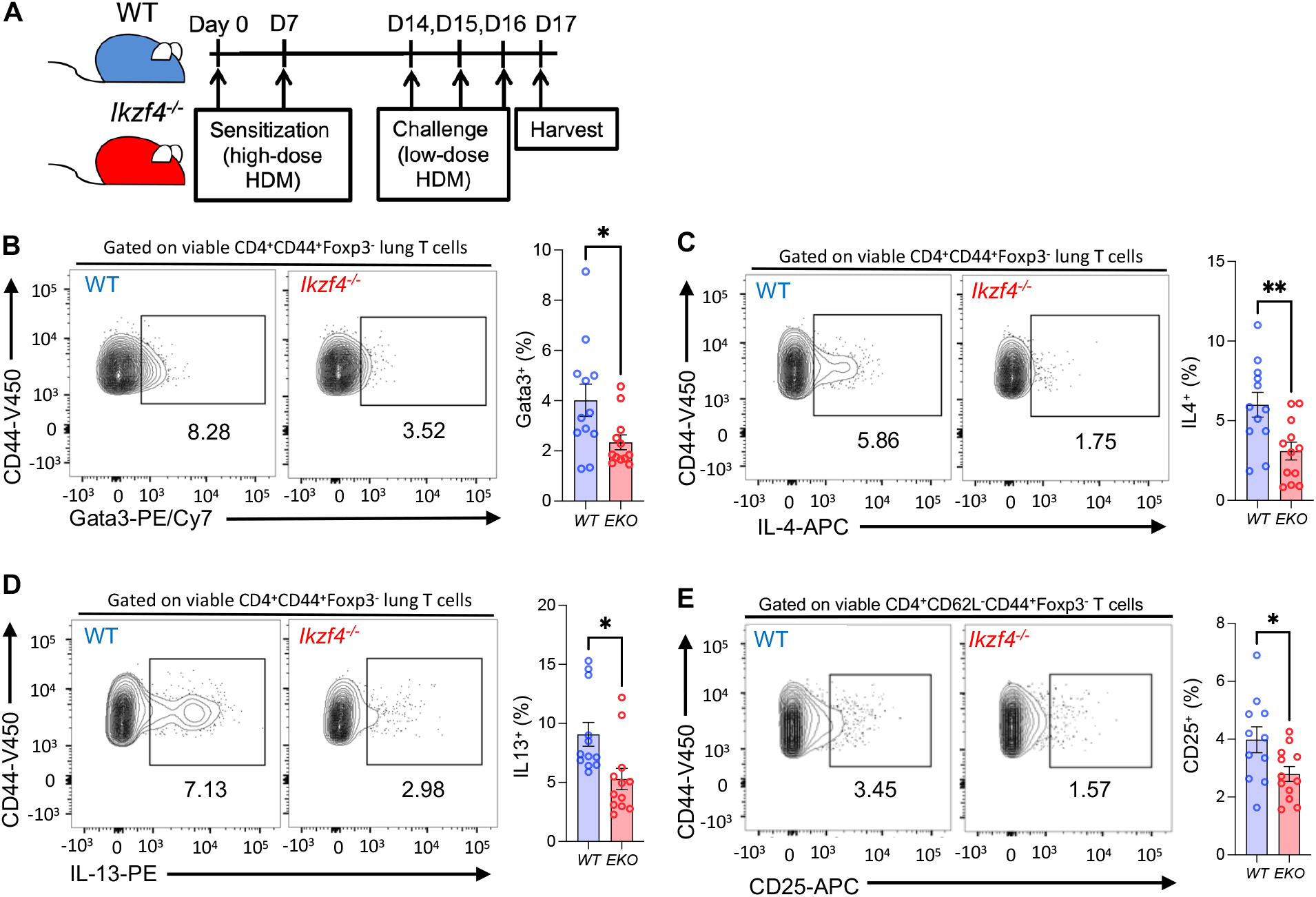
Eos-deficient mice have defective T_H_2 induction and effector responses. **(A)** WT or Eos-deficient (*Ikzf4^-/-^*) C57BL/6J mice were sensitized on days 0 and 7 with 10 μg of house dust mite (HDM) solution via oropharyngeal (o.p.) aspiration. On days 14, 15, and 16, mice were challenged o.p. with 2 μg of the HDM preparation. On day 17, lungs were harvested and viable CD4^+^ effector populations were analyzed via flow cytometry. **(B-D)** Single-cell suspensions from the lung were incubated in culture medium in the presence of PMA and ionomycin stimulation and protein transport inhibitors for 4 hours. Gata3 protein expression and IL-4 and IL-13 production of viable CD4^+^CD44^+^Foxp3^-^ populations was analyzed via flow cytometry. Data are compiled from 3 independent experiments (n = 12 ± s.e.m; \**P* < 0.05, \*\**P* < 0.01; unpaired Student’s *t*-test). **(E-G)** CD25 (IL-2Ra) surface expression was evaluated on viable CD4^+^CD44^+^CD62L^-^Foxp3^-^ (effector) T cells from the lung via flow cytometry. Percentage of CD25^+^ cells (E) and median fluorescence intensity (MFI) for Foxp3^-^ (F) and Foxp3^-^CD25^+^ (G) cell populations are shown. Data are representative of 3 independent experiments. (n = 11-12 ± s.e.m; *\*P* < 0.05, *\*\*P* < 0.01; unpaired Student’s *t*-test).

Compared to their WT counterparts, the Eos-deficient mice had significantly lower levels of CD4^+^CD62L^-^CD44^+^Foxp3^-^Gata3^+^ cells, as assessed by percent positive cells (**Fig. 4B**). Additionally, of those CD4^+^CD62L^-^CD44^+^Foxp3^-^ T cells, Eos-deficient cells produced significantly lower levels of IL-4 and IL-13 (**Fig. 4C,D**). We also considered that Eos deficiency may cause a decrease in immunosuppressive T_REG_ cells, enabling uncontrolled T_H_2 proliferation. However, our data did not show a drop in the number of CD4^+^CD62L^-^CD44^+^Foxp3^+^ T cells present in the context of global Eos deficiency (**Supplemental Fig. 3A**). We further analyzed protein expression of CD25 in CD4^+^CD62L^-^CD44^+^Foxp3^-^ T_H_2 cells by flow cytometry and found that a lack of Eos resulted in significant decreases in the number of CD25 percent positive cells (**Fig. 4E)**. This decrease in CD25 with Eos deficiency was also reflected in the percent positive populations of the CD4^+^CD62L^-^CD44^+^Foxp3^+^ T_REG_ population (**Supplemental Fig. 3B**), further supporting at least a partially conserved mechanism for Eos and the IL-2/STAT5 pathway across T cell subsets. Overall, these data show that in an in vivo house dust mite allergic asthma model, Eos-deficient mice have a lower induction of T_H_2 differentiation via downregulated Gata3 and IL-2R and effector cytokine production.

### 5. *Eos and STAT5 form a transcriptional complex that upregulates STAT5 tyrosine phosphorylation and promotes robust* Il4ra *expression*

A key finding from our prior studies was that Aiolos interacts with STAT3 to exert transcriptional changes at the Bcl-6 promoter (36). Because of the high degree of homology among IkZF family members and STAT proteins, we hypothesized that Eos may be impacting T_H_2 differentiation and effector cytokine production via direct interaction with STAT5. We performed co-immunoprecipitation (Co-IP) assays and identified a novel Eos-STAT5 complex in in vitro-generated T_H_2 cells (**Fig. 5A**), as well as in in vitro T_REG_ cells (**Supplemental Fig. 3C**). To support these findings, we constructed vectors for Eos and STAT5B^CA^ and overexpressed them in murine EL4 T cell thymoma cell line to identify this Eos-STAT5 complex (**Fig. 5B**). We found that this interaction with STAT5B^CA^ was specific to Eos, as other IkZF members Ikaros or Aiolos did not co-immunoprecipitate with STAT5 (**Fig. 5C**). We then wanted to determine the relevance of this Eos-STAT5 complex in T_H_2 differentiation. Previous literature has defined how STAT5 binding sites induces CD124 (IL-4Rα) expression (18), which is critical for robust T_H_2 differentiation; indeed, we found that CD124 was downregulated in our in vitro T_H_2 transcriptomic analysis (**Fig. 2C**). To assess how the Eos-STAT5 complex may contribute to *Il4ra* expression, we used an EL4 overexpression system and found that *Il4ra* expression was highest when Eos^WT^ and STAT5B^CA^ were co-expressed compared to either protein overexpressed alone (**Fig. 5D**). Intriguingly, we found that when Eos^WT^ and STAT5B^CA^ were overexpressed together, there was a marked increase in the amount of pY-STAT5 protein on immunoblot compared to either protein overexpressed alone (**Fig. 5E**), suggesting this Eos-STAT5 interaction bolsters pY-STAT5 levels. Because this system uses a constitutively active STAT5 and EL4 T cells do not respond to environmental IL-2 signals, these results suggest that STAT5 tyrosine phosphorylation is dependent on interactions with Eos rather than environmental cytokines. Further supporting this, we found that overexpression of Eos, but not Ikaros or Aiolos, resulted in increased pY-STAT5 (**Fig. 5F**). Collectively, these data indicate an Eos and STAT5 interaction correlates with increased gene expression of STAT5 target genes and increased STAT5 tyrosine phosphorylation.

**Figure 5.**
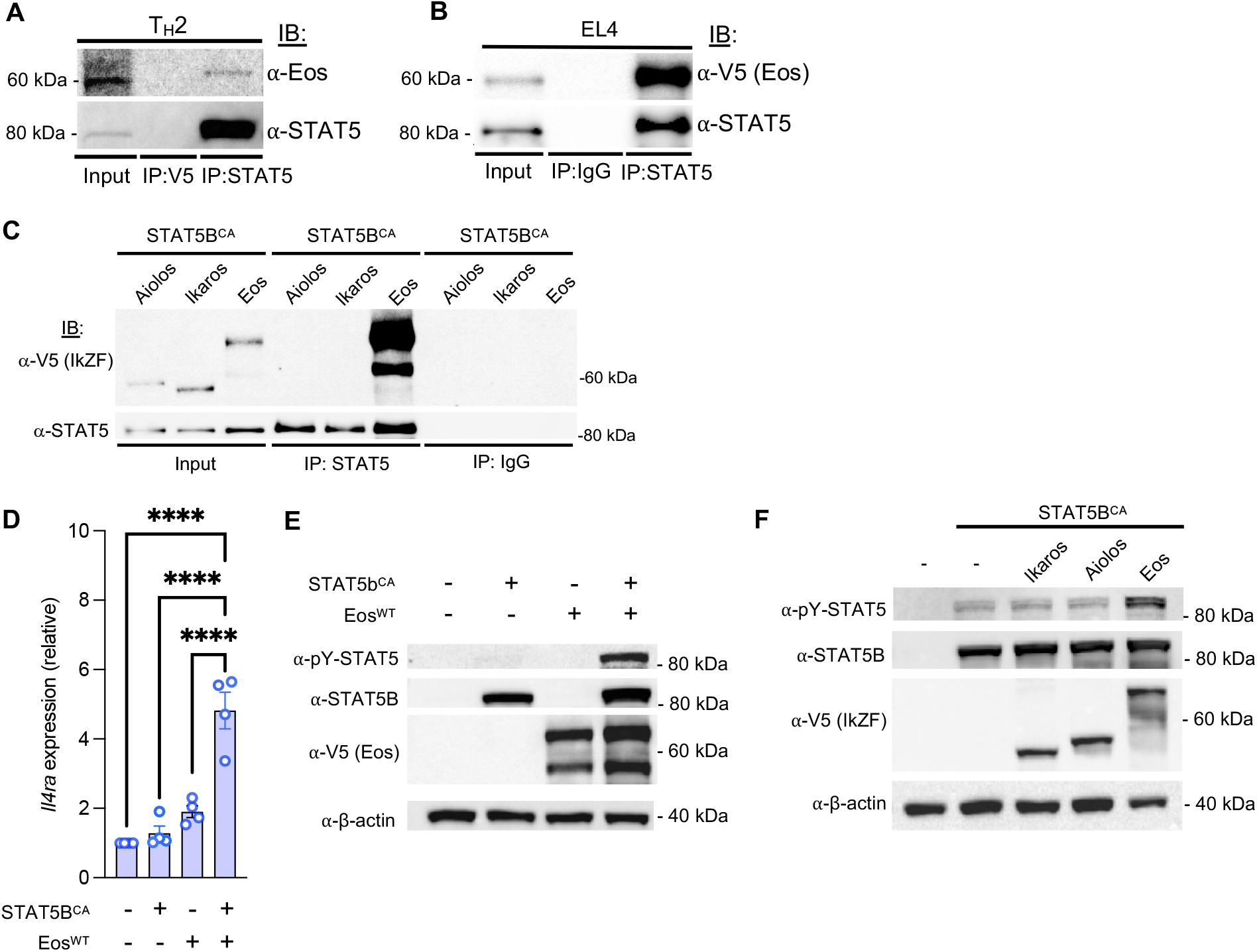
Eos and STAT5 form a transcription factor complex and cooperate to regulate IL-4Rα. **(A)** Co-Immunoprecipitation (Co-IP) of endogenously expressed proteins in T_H_2 cells with control Ab (α-V5) or α-STAT5, followed by immunoblot with anti-Eos (n = 3 independent experiments). **(B)** Co-IP analysis of overexpressed V5-tagged Eos and tagless STAT5B^CA^ in EL4 T cells. Lysates were immunoprecipitated with anti-STAT5, followed by immunoblot analysis with α-V5 (for detecting Eos) (n = 3 independent experiments). **(C)** Co-IP analysis of overexpressed Aiolos, Ikaros, or Eos with tagless STAT5B^CA^ in EL4 T cells. Lysates were immunoprecipitated with α-STAT5, followed by immunoblot analysis with α-V5 (for detecting IkZF factors) (n = 3 independent experiments). **(D)** RNA was isolated from EL4 cells transfected with STAT5B^CA^, Eos, or STAT5B^CA^ and Eos in combination. qRT-PCR was used to assess *Il4ra* expression. Data were normalized to *Rps18* and presented as fold change relative to the empty vector sample (n = 4 independent experiments, mean ± s.e.m., \*\*\*\**P* < 0.0001; one-way ANOVA with Tukey’s post-hoc test). **(E)** Immunoblot analysis of pY-STAT5, STAT5, and Eos protein expression in EL4 T cells transfected with STAT5B^CA^, Eos, or STAT5B^CA^ and Eos in combination. β-actin serves as a loading control. Shown is a representative blot of 3 independent experiments. **(F)** Immunoblot analysis of pY-STAT5 and STAT5 in EL4 T cells transfected with tagless STAT5B^CA^ alone or in combination with the indicated IkZF. β-actin serves as a loading control (n = 3 independent experiments).

### 6. The C-terminal domain of Eos is required for binding to and tyrosine-phosphorylated activation of STAT5

To parse out how Eos interacts with STAT5, we created deletion mutants of the conserved N-terminal DNA-binding (Eos^ΔZF1,2^) and C-terminal protein-protein interaction (Eos^ΔC^) domains (**Fig. 6A**). Overexpression of these Eos mutants with STAT5B^CA^ and subsequent Co-IP revealed that Eos^ΔC^, but not Eos^ΔZF1,2^, lost the ability to interact with STAT5 (**Fig. 6B**). We also wanted to determine whether the Eos-STAT5 interaction was required for STAT5 tyrosine phosphorylation. When we overexpressed the mutant Eos vectors with STAT5B^CA^, we observed that Eos^ΔC^ co-expressed with STAT5B^CA^ had lower pY-STAT5 levels than either Eos^WT^ overexpressed with STAT5B^CA^ or Eos^ΔZF1,2^ overexpressed with STAT5B^CA^ (**Fig. 6C**), suggesting that a robust Eos-STAT5 interaction is required to increase pY-STAT5 levels.

**Figure 6:**
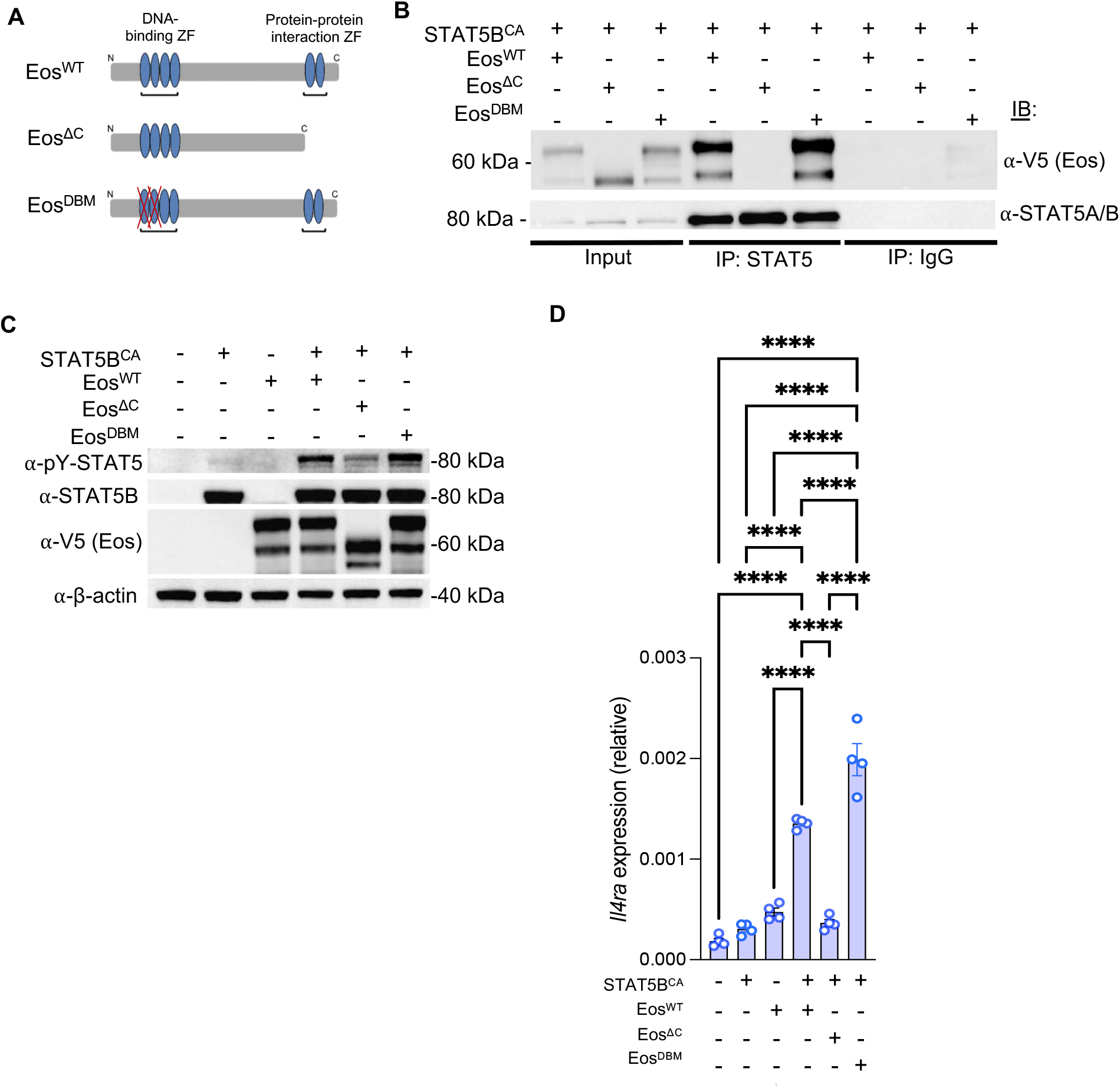
The C-terminal domain of Eos is required for binding to and activating STAT5. **(A)** Deletion mutants of the conserved N-terminal DNA-binding (Eos^ΔZF1,2^) and C-terminal protein-protein interaction (Eos^ΔC^) domains were created using the Agilent QuikChange Site-Directed Mutagenesis Kit. Wildtype and mutant coding sequences were transferred to the pEF1/V5-His vector for overexpression in EL4 T cells. **(B)** Co-IP of V5-tagged wild-type Eos (Eos^WT^), Eos^ΔC^, and Eos^ΔZF1,2^ overexpressed with tagless STAT5B^CA^ in EL4 T cells. Lysates were immunoprecipitated with anti-STAT5, followed by immunoblot analysis with α-V5 for detecting Eos proteins (n = 3 independent experiments). **(C)** Immunoblot analysis of pY-STAT5, STAT5, and Eos protein expression in EL4 T cells transfected with tagless STAT5B^CA^ and either V5-tagged Eos^WT^, Eos^ΔC^, or Eos^ΔZF1,2^. β-actin serves as a loading control (n = 3 independent experiments). **(D)** EL4 T cells were transfected with tagless STAT5B^CA^ and either V5-tagged Eos^WT^, Eos^ΔC^, or Eos^ΔZF1,2^. After 22h, RNA was isolated, and *Il4ra* expression was measured by qRT-PCR. Data were normalized to *Rps18* as a control and presented as fold change in expression relative to *Rps18*. (n = 4 independent experiments, mean ± s.e.m., ****P < 0.0001).

We then wanted to establish whether Eos and the IL-2/STAT5 axis work together to regulate T_H_2 genes. Because CD25 is known to positively regulate the expression of CD124 (18), we tested how the absence of the Eos-STAT5 interaction may impact downstream regulation of CD124 expression. Transcript levels of *Il4ra* showed significant decreases when Eos^ΔC^ was overexpressed with STAT5B^CA^ compared to Eos^WT^ or Eos^ΔZF1,2^ overexpressed with STAT5B^CA^ (**Fig. 6D**). Overall, these data show that in EL4 T cells, Eos and STAT5 cooperate to regulate the expression of *Il4ra* to specifically control T_H_2 differentiation.

## Discussion

For roughly a decade, Eos has been categorized as a transcription factor important for maintaining both CD4^+^ T_REG_ cell identity and associated suppressive functions (46–49, 66, 67). More recent work has suggested that Eos plays additional roles in regulating the differentiation and function of effector T cell populations (46, 50). However, the precise role of Eos in these processes, which are essential for mammalian adaptive immunity, has remained elusive. To this end, we now demonstrate that Eos is a novel regulator of T_H_2 cell differentiation and effector cytokine production. We find that Eos-deficient mice have reduced T_H_2 induction in a murine house dust mite asthma model, including defects to T_H_2 differentiation and diminished effector cytokine production, as measured by expression of Gata3 and production of T_H_2 cytokines. Whole-transcriptome analysis revealed global defects in the expression of the T_H_2 gene program in Eos-deficient CD4^+^ T cells. Mechanistically, we observed a substantial decrease in the activation of STAT5 in the absence of Eos. Thus, our findings implicate Eos as an important regulator of T_H_2 differentiation and suggest that augmentation of STAT5 activity represents at least one mechanism by which Eos promotes the T_H_2 gene program.

Previous work from our laboratory demonstrated that the IkZF family member Aiolos physically interacts and cooperates with STAT3 to positively regulate Bcl-6 expression in CD4^+^ T cell populations (36). Here, we show that Eos and STAT5 interact in T_H_2 cells, and disruption of this interaction at the C-terminal ZF domain of Eos, which is known to mediate homo- and hetero-dimerization amongst IkZF transcription factors, dysregulates T_H_2 gene expression patterns (28, 68). This is consistent with our previous findings for Aiolos-STAT3 interactions, establishing the C-terminal ZF domain of Ikaros family members as a conserved regulatory feature that enables the formation of IkZF-STAT factor regulatory modules (36). Our data also demonstrate that Eos^ΔC^ overexpressed with STAT5B^CA^ still has greater pY-STAT5 levels than wild-type Eos (Eos^WT^) expressed alone, prompting further discussion of other IkZF/STAT5 interacting domains beyond the IkZF C-terminal ZF domain that may regulate STAT activity and function. Furthermore, we show that distinct cytokine signatures drive the formation and function of the Eos/STAT5 and Aiolos/STAT3 modules in T helper cell populations (36). Thus, it is intriguing to speculate that additional cytokine signals driving T_H_2 differentiation (e.g. IL-4/STAT6), as well as those cytokine-STAT pathways that direct differentiation of other T helper subsets, may promote the formation and function of distinct IkZF-STAT regulatory complexes.

The precise mechanisms by which Eos regulates target gene expression are incompletely defined. Current evidence indicates that IkZF factors primarily recruit remodeling complexes to alter chromatin structure (28, 29, 69). However, our data in T_H_2 cells point to a novel mechanism whereby Eos works in a feed-forward fashion to promote STAT5 activation and subsequent association with target genes. While how Eos modulates STAT5 phosphorylation is currently unclear, we hypothesize that there are several non-mutually exclusive possibilities. First, and most clearly, our data indicate that Eos regulates expression of IL-2 receptor subunits, through which STAT5 is activated in the T_H_2 cell population. Indeed, a prior study described Eos-dependent regulation of CD25 in undifferentiated, activated CD4^+^ T_CONV_ cell populations (50). While much of the data we present in T_H_2 cells are consistent with this mechanism, it is important to note that our overexpression studies were performed with a constitutively active STAT5B—the dominant STAT5 isoform in effector and regulatory T cell responses (62)—in the absence of signals from IL-2. Thus, we speculate that Eos’ modulation of STAT5 activity may result from altered expression and/or function of kinases or phosphatases that act upon STAT5. This includes the possibility that the physical interaction between Eos and STAT5 enhances STAT5 protein stability and/or protects it from phosphatase activity. Ultimately, future studies will be necessary to comprehensively evaluate the mechanism(s) by which Eos influences STAT5 phosphorylation.

Our findings also offer the opportunity to compare roles for Eos in T_REG_ and effector T_H_2 cell populations. In T_REG_ cells, Eos functions as a transcriptional repressor that works in concert with Foxp3 to direct T_REG_ cell-specific expression patterns (47, 48, 66). Notably, a known Eos target in T_REG_ cells is the *Il2* locus (48). However, findings from our study suggest that Eos-dependent repression of IL-2 is not conserved in T_H_2 cells. While we cannot definitively rule out a repressive role for Eos in T_H_2 cells, our findings suggest that Eos plays an important role in positively regulating the T_H_2 gene program by potentiating STAT5 activity. Curiously, Eos is expressed at much higher levels in T_REG_ versus effector T cell populations (50). As such, our findings suggest that Eos expression may operate as a rheostat controlling the amplitude of the IL-2/STAT5 signaling pathway in different T helper cell subsets. Functionally, this may be particularly advantageous for T_REG_ cells, as it may provide a mechanism to maintain high levels of active STAT5 in low IL-2 environments (70, 71). This novel Eos-STAT5 relationship also is likely to be clinically relevant, as therapies employing low doses of IL-2 have been used in efforts to preferentially enhance T_REG_ cell populations for the treatment of a variety of autoimmune disorders (72, 73). A comprehensive understanding of the conserved and/or divergent roles for Eos in driving the transcriptional programs of T_REG_ and T_H_2 cell populations awaits additional study, including those that will probe other CD4^+^ T cell subsets dependent on IL-2 signaling for their differentiation (e.g. T_H_1 or T_H_9 cells) (18, 74–76).

Finally, it will be important for future studies to assess whether Eos-STAT5 regulatory modules function downstream of cytokine signals other than IL-2. Indeed, STAT5 is also activated downstream of signals from the cytokines IL-7 and IL-15, which impart diverse functions across several immune cell populations (77, 78). As such, it will be of interest to assess potential cooperative roles for Eos and STAT5 in CD8^+^ T, B, natural killer, and innate lymphoid cell populations (79). Ultimately, these studies will generate important insights into general mechanisms by which Ikaros family members regulate the differentiation and function of immune cell populations, which may be leveraged in targeted immunotherapy approaches to treat human disease.

## Acknowledgments

The authors would like to thank all members of the Oestreich Lab, as well as colleagues in the Department of Microbial Infection and Immunity for constructive feedback.

## Author contributions

J.A.T. assisted with the design of the study, performed experiments, analyzed data, and wrote the manuscript. K.A.R. and B.K.S. assisted with the design of the study, performed experiments, and analyzed data. D.M.J., R.T.W., M.D.P., M.N.R., E.G.D., and L.M.C. performed experiments and analyzed data. M.J.Y., S.V., and K.M.G. assisted with the house dust mite infection experiments. K.J.O. supervised the research, designed the study, analyzed data, and wrote the manuscript.

## Data and materials availability

RNA-seq data sets have been deposited in the GEO repository under accession number GSE216737. All other datasets and materials from this study will be made available upon reasonable request. Requests should be sent to the corresponding author.

## References

1. Coffman, R. L., B. W. Seymour, S. Hudak, J. Jackson, and D. Rennick. 1989. Antibody to interleukin-5 inhibits helminth-induced eosinophilia in mice. Science 245: 308–310.

2. Grzych, J. M., E. Pearce, A. Cheever, Z. A. Caulada, P. Caspar, S. Heiny, F. Lewis, and A. Sher. 1991. Egg deposition is the major stimulus for the production of Th2 cytokines in murine schistosomiasis mansoni. J Immunol 146: 1322–1327.

3. Allen, J. E., and T. E. Sutherland. 2014. Host protective roles of type 2 immunity: parasite killing and tissue repair, flip sides of the same coin. Semin Immunol 26: 329–340.

4. Hall, S., and D. K. Agrawal. 2014. Key mediators in the immunopathogenesis of allergic asthma. Int Immunopharmacol 23: 316–329.

5. Vale, K. 2016. Targeting the JAK-STAT pathway in the treatment of ‘Th2-high’ severe asthma. Future Med Chem 8: 405–419.

6. O’Shea, J. J., and W. E. Paul. 2010. Mechanisms underlying lineage commitment and plasticity of helper CD4+ T cells. Science 327: 1098–1102.

7. Bonelli, M., H. Y. Shih, K. Hirahara, K. Singelton, A. Laurence, A. Poholek, T. Hand, Y. Mikami, G. Vahedi, Y. Kanno, and J. J. O’Shea. 2014. Helper T cell plasticity: impact of extrinsic and intrinsic signals on transcriptomes and epigenomes. Curr Top Microbiol Immunol 381: 279–326.

8. Maier, E., A. Duschl, and J. Horejs-Hoeck. 2012. STAT6-dependent and-independent mechanisms in Th2 polarization. Eur J Immunol 42: 2827–2833.

9. Shimoda, K., J. van Deursen, M. Y. Sangster, S. R. Sarawar, R. T. Carson, R. A. Tripp, C. Chu, F. W. Quelle, T. Nosaka, D. A. Vignali, P. C. Doherty, G. Grosveld, W. E. Paul, and J. N. Ihle. 1996. Lack of IL-4-induced Th2 response and IgE class switching in mice with disrupted Stat6 gene. Nature 380: 630–633.

10. Takeda, K., T. Tanaka, W. Shi, M. Matsumoto, M. Minami, S. Kashiwamura, K. Nakanishi, N. Yoshida, T. Kishimoto, and S. Akira. 1996. Essential role of Stat6 in IL-4 signalling. Nature 380: 627–630.

11. Jankovic, D., M. C. Kullberg, N. Noben-Trauth, P. Caspar, W. E. Paul, and A. Sher. 2000. Single cell analysis reveals that IL-4 receptor/Stat6 signaling is not required for the in vivo or in vitro development of CD4+ lymphocytes with a Th2 cytokine profile. J Immunol 164: 3047–3055.

12. Cote-Sierra, J., G. Foucras, L. Guo, L. Chiodetti, H. A. Young, J. Hu-Li, J. Zhu, and W. E. Paul. 2004. Interleukin 2 plays a central role in Th2 differentiation. Proc Natl Acad Sci U S A 101: 3880–3885.

13. Le Gros, G., S. Z. Ben-Sasson, R. Seder, F. D. Finkelman, and W. E. Paul. 1990. Generation of interleukin 4 (IL-4)-producing cells in vivo and in vitro: IL-2 and IL-4 are required for in vitro generation of IL-4-producing cells. J Exp Med 172: 921–929.

14. Zhu, J., J. Cote-Sierra, L. Guo, and W. E. Paul. 2003. Stat5 activation plays a critical role in Th2 differentiation. Immunity 19: 739–748.

15. Zhu, J. 2015. T helper 2 (Th2) cell differentiation, type 2 innate lymphoid cell (ILC2) development and regulation of interleukin-4 (IL-4) and IL-13 production. Cytokine 75: 14–24.

16. Ross, S. H., and D. A. Cantrell. 2018. Signaling and Function of Interleukin-2 in T Lymphocytes. Annu Rev Immunol 36: 411–433.

17. Walker, J. A., and A. N. J. McKenzie. 2018. TH2 cell development and function. Nat Rev Immunol 18: 121–133.

18. Liao, W., D. E. Schones, J. Oh, Y. Cui, K. Cui, T. Y. Roh, K. Zhao, and W. J. Leonard. 2008. Priming for T helper type 2 differentiation by interleukin 2-mediated induction of interleukin 4 receptor alpha-chain expression. Nat Immunol 9: 1288–1296.

19. Lin, J. X., and W. J. Leonard. 2000. The role of Stat5a and Stat5b in signaling by IL-2 family cytokines. Oncogene 19: 2566–2576.

20. Guo, L., G. Wei, J. Zhu, W. Liao, W. J. Leonard, K. Zhao, and W. Paul. 2009. IL-1 family members and STAT activators induce cytokine production by Th2, Th17, and Th1 cells. Proc Natl Acad Sci U S A 106: 13463–13468.

21. Laurence, A., C. M. Tato, T. S. Davidson, Y. Kanno, Z. Chen, Z. Yao, R. B. Blank, F. Meylan, R. Siegel, L. Hennighausen, E. M. Shevach, and J. O’Shea J. 2007. Interleukin-2 signaling via STAT5 constrains T helper 17 cell generation. Immunity 26: 371–381.

22. Ballesteros-Tato, A., B. Leon, B. A. Graf, A. Moquin, P. S. Adams, F. E. Lund, and T. D. Randall. 2012. Interleukin-2 inhibits germinal center formation by limiting T follicular helper cell differentiation. Immunity 36: 847–856.

23. Johnston, R. J., Y. S. Choi, J. A. Diamond, J. A. Yang, and S. Crotty. 2012. STAT5 is a potent negative regulator of TFH cell differentiation. J Exp Med 209: 243–250.

24. Johnston, R. J., A. C. Poholek, D. DiToro, I. Yusuf, D. Eto, B. Barnett, A. L. Dent, J. Craft, and S. Crotty. 2009. Bcl6 and Blimp-1 are reciprocal and antagonistic regulators of T follicular helper cell differentiation. Science 325: 1006–1010.

25. Kallies, A., E. D. Hawkins, G. T. Belz, D. Metcalf, M. Hommel, L. M. Corcoran, P. D. Hodgkin, and S. L. Nutt. 2006. Transcriptional repressor Blimp-1 is essential for T cell homeostasis and self-tolerance. Nat Immunol 7: 466–474.

26. Fu, S. H., L. T. Yeh, C. C. Chu, B. L. Yen, and H. K. Sytwu. 2017. New insights into Blimp-1 in T lymphocytes: a divergent regulator of cell destiny and effector function. J Biomed Sci 24: 49.

27. He, K., A. Hettinga, S. L. Kale, S. Hu, M. M. Xie, A. L. Dent, A. Ray, and A. C. Poholek. 2020. Blimp-1 is essential for allergen-induced asthma and Th2 cell development in the lung. J Exp Med 217.

28. Yoshida, T., and K. Georgopoulos. 2014. Ikaros fingers on lymphocyte differentiation. Int J Hematol 100: 220–229.

29. Heizmann, B., P. Kastner, and S. Chan. 2018. The Ikaros family in lymphocyte development. Curr Opin Immunol 51: 14–23.

30. Quirion, M. R., G. D. Gregory, S. E. Umetsu, S. Winandy, and M. A. Brown. 2009. Cutting edge: Ikaros is a regulator of Th2 cell differentiation. J Immunol 182: 741–745.

31. Hoshino, A., D. Boutboul, Y. Zhang, H. S. Kuehn, J. Hadjadj, N. Ozdemir, T. Celkan, C. Walz, C. Picard, C. Lenoir, N. Mahlaoui, C. Klein, X. Peng, A. Azar, E. Reigh, M. Cheminant, A. Fischer, F. Rieux-Laucat, I. Callebaut, F. Hauck, J. Milner, S. D. Rosenzweig, and S. Latour. 2022. Gain-of-function IKZF1 variants in humans cause immune dysregulation associated with abnormal T/B cell late differentiation. Sci Immunol 7: eabi7160.

32. Wong, L. Y., J. K. Hatfield, and M. A. Brown. 2013. Ikaros sets the potential for Th17 lineage gene expression through effects on chromatin state in early T cell development. J Biol Chem 288: 35170–35179.

33. Lyon de Ana, C., K. Arakcheeva, P. Agnihotri, N. Derosia, and S. Winandy. 2019. Lack of Ikaros Deregulates Inflammatory Gene Programs in T Cells. J Immunol 202: 1112–1123.

34. Bernardi, C., G. Maurer, T. Ye, P. Marchal, B. Jost, M. Wissler, U. Maurer, P. Kastner, S. Chan, and C. Charvet. 2021. CD4(+) T cells require Ikaros to inhibit their differentiation toward a pathogenic cell fate. Proc Natl Acad Sci U S A 118.

35. Thomas, R. M., N. Chunder, C. Chen, S. E. Umetsu, S. Winandy, and A. D. Wells. 2007. Ikaros enforces the costimulatory requirement for IL2 gene expression and is required for anergy induction in CD4+ T lymphocytes. J Immunol 179: 7305–7315.

36. Read, K. A., M. D. Powell, C. E. Baker, B. K. Sreekumar, V. M. Ringel-Scaia, H. Bachus, R. E. Martin, I. D. Cooley, I. C. Allen, A. Ballesteros-Tato, and K. J. Oestreich. 2017. Integrated STAT3 and Ikaros Zinc Finger Transcription Factor Activities Regulate Bcl-6 Expression in CD4(+) Th Cells. J Immunol 199: 2377–2387.

37. Heller, J. J., H. Schjerven, S. Li, A. Lee, J. Qiu, Z. M. Chen, S. T. Smale, and L. Zhou. 2014. Restriction of IL-22-producing T cell responses and differential regulation of regulatory T cell compartments by zinc finger transcription factor Ikaros. J Immunol 193: 3934–3946.

38. Agnihotri, P., N. M. Robertson, S. E. Umetsu, K. Arakcheeva, and S. Winandy. 2017. Lack of Ikaros cripples expression of Foxo1 and its targets in naive T cells. Immunology 152: 494–506.

39. Baine, I., S. Basu, R. Ames, R. S. Sellers, and F. Macian. 2013. Helios induces epigenetic silencing of IL2 gene expression in regulatory T cells. J Immunol 190: 1008–1016.

40. Kim, H. J., R. A. Barnitz, T. Kreslavsky, F. D. Brown, H. Moffett, M. E. Lemieux, Y. Kaygusuz, T. Meissner, T. A. Holderried, S. Chan, P. Kastner, W. N. Haining, and H. Cantor. 2015. Stable inhibitory activity of regulatory T cells requires the transcription factor Helios. Science 350: 334–339.

41. Raffin, C., P. Pignon, C. Celse, E. Debien, D. Valmori, and M. Ayyoub. 2013. Human memory Helios-FOXP3+ regulatory T cells (Tregs) encompass induced Tregs that express Aiolos and respond to IL-1beta by downregulating their suppressor functions. J Immunol 191: 4619–4627.

42. Serre, K., C. Benezech, G. Desanti, S. Bobat, K. M. Toellner, R. Bird, S. Chan, P. Kastner, A. F. Cunningham, I. C. Maclennan, and E. Mohr. 2011. Helios is associated with CD4 T cells differentiating to T helper 2 and follicular helper T cells in vivo independently of Foxp3 expression. PLoS One 6: e20731.

43. Quintana, F. J., H. Jin, E. J. Burns, M. Nadeau, A. Yeste, D. Kumar, M. Rangachari, C. Zhu, S. Xiao, J. Seavitt, K. Georgopoulos, and V. K. Kuchroo. 2012. Aiolos promotes TH17 differentiation by directly silencing Il2 expression. Nat Immunol 13: 770–777.

44. Fujimura, K., A. Oyamada, Y. Iwamoto, Y. Yoshikai, and H. Yamada. 2013. CD4 T cell-intrinsic IL-2 signaling differentially affects Th1 and Th17 development. J Leukoc Biol 94: 271–279.

45. Lim, H. W., S. G. Kang, J. K. Ryu, B. Schilling, M. Fei, I. S. Lee, A. Kehasse, K. Shirakawa, M. Yokoyama, M. Schnolzer, H. G. Kasler, H. S. Kwon, B. W. Gibson, H. Sato, K. Akassoglou, C. Xiao, D. R. Littman, M. Ott, and E. Verdin. 2015. SIRT1 deacetylates RORgammat and enhances Th17 cell generation. J Exp Med 212: 607–617.

46. Gokhale, A. S., A. Gangaplara, M. Lopez-Occasio, A. M. Thornton, and E. M. Shevach. 2019. Selective deletion of Eos (Ikzf4) in T-regulatory cells leads to loss of suppressive function and development of systemic autoimmunity. J Autoimmun 105: 102300.

47. Sharma, M. D., L. Huang, J. H. Choi, E. J. Lee, J. M. Wilson, H. Lemos, F. Pan, B. R. Blazar, D. M. Pardoll, A. L. Mellor, H. Shi, and D. H. Munn. 2013. An inherently bifunctional subset of Foxp3+ T helper cells is controlled by the transcription factor eos. Immunity 38: 998–1012.

48. Pan, F., H. Yu, E. V. Dang, J. Barbi, X. Pan, J. F. Grosso, D. Jinasena, S. M. Sharma, E. M. McCadden, D. Getnet, C. G. Drake, J. O. Liu, M. C. Ostrowski, and D. M. Pardoll. 2009. Eos mediates Foxp3-dependent gene silencing in CD4+ regulatory T cells. Science 325: 1142–1146.

49. Yang, H. Y., J. Barbi, C. Y. Wu, Y. Zheng, P. D. Vignali, X. Wu, J. H. Tao, B. V. Park, S. Bandara, L. Novack, X. Ni, X. Yang, K. Y. Chang, R. C. Wu, J. Zhang, C. W. Yang, D. M. Pardoll, H. Li, and F. Pan. 2016. MicroRNA-17 Modulates Regulatory T Cell Function by Targeting Co-regulators of the Foxp3 Transcription Factor. Immunity 45: 83–93.

50. Rieder, S. A., A. Metidji, D. D. Glass, A. M. Thornton, T. Ikeda, B. A. Morgan, and E. M. Shevach. 2015. Eos Is Redundant for Regulatory T Cell Function but Plays an Important Role in IL-2 and Th17 Production by CD4+ Conventional T Cells. J Immunol 195: 553–563.

51. Liu, S. Q., S. Jiang, C. Li, B. Zhang, and Q. J. Li. 2014. miR-17-92 cluster targets phosphatase and tensin homology and Ikaros Family Zinc Finger 4 to promote TH17-mediated inflammation. J Biol Chem 289: 12446–12456.

52. Yung, J. A., H. Fuseini, and D. C. Newcomb. 2018. Hormones, sex, and asthma. Ann Allergy Asthma Immunol 120: 488–494.

53. Cheng, C., H. Wu, M. Wang, L. Wang, H. Zou, S. Li, and R. Liu. 2019. Estrogen ameliorates allergic airway inflammation by regulating activation of NLRP3 in mice. Biosci Rep 39.

54. Kalidhindi, R. S. R., N. S. Ambhore, S. Bhallamudi, J. Loganathan, and V. Sathish. 2019. Role of Estrogen Receptors alpha and beta in a Murine Model of Asthma: Exacerbated Airway Hyperresponsiveness and Remodeling in ERbeta Knockout Mice. Front Pharmacol 10: 1499.

55. Farrar, M. A. 2010. Design and use of constitutively active STAT5 constructs. Methods Enzymol 485: 583–596.

56. Oestreich, K. J., S. E. Mohn, and A. S. Weinmann. 2012. Molecular mechanisms that control the expression and activity of Bcl-6 in TH1 cells to regulate flexibility with a TFH-like gene profile. Nat Immunol 13: 405–411.

57. Oestreich, K. J., K. A. Read, S. E. Gilbertson, K. P. Hough, P. W. McDonald, V. Krishnamoorthy, and A. S. Weinmann. 2014. Bcl-6 directly represses the gene program of the glycolysis pathway. Nat Immunol 15: 957–964.

58. Goedhart, J., and M. S. Luijsterburg. 2020. VolcaNoseR is a web app for creating, exploring, labeling and sharing volcano plots. Sci Rep 10: 20560.

59. McDonald, P. W., K. A. Read, C. E. Baker, A. E. Anderson, M. D. Powell, A. Ballesteros-Tato, and K. J. Oestreich. 2016. IL-7 signalling represses Bcl-6 and the TFH gene program. Nat Commun 7: 10285.

60. Thomas, R. M., C. Chen, N. Chunder, L. Ma, J. Taylor, E. J. Pearce, and A. D. Wells. 2010. Ikaros silences T-bet expression and interferon-gamma production during T helper 2 differentiation. J Biol Chem 285: 2545–2553.

61. Jones, D. M., K. A. Read, and K. J. Oestreich. 2020. Dynamic Roles for IL-2-STAT5 Signaling in Effector and Regulatory CD4(+) T Cell Populations. J Immunol 205: 1721–1730.

62. Villarino, A., A. Laurence, G. W. Robinson, M. Bonelli, B. Dema, B. Afzali, H. Y. Shih, H. W. Sun, S. R. Brooks, L. Hennighausen, Y. Kanno, and J. J. O’Shea. 2016. Signal transducer and activator of transcription 5 (STAT5) paralog dose governs T cell effector and regulatory functions. Elife 5.

63. Able, A. A., J. A. Burrell, and J. M. Stephens. 2017. STAT5-Interacting Proteins: A Synopsis of Proteins that Regulate STAT5 Activity. Biology (Basel) 6.

64. Gouilleux, F., H. Wakao, M. Mundt, and B. Groner. 1994. Prolactin induces phosphorylation of Tyr694 of Stat5 (MGF), a prerequisite for DNA binding and induction of transcription. EMBO J 13: 4361–4369.

65. Whitehead, G. S., S. Y. Thomas, and D. N. Cook. 2014. Modulation of distinct asthmatic phenotypes in mice by dose-dependent inhalation of microbial products. Environ Health Perspect 122: 34–42.

66. Bettini, M. L., F. Pan, M. Bettini, D. Finkelstein, J. E. Rehg, S. Floess, B. D. Bell, S. F. Ziegler, J. Huehn, D. M. Pardoll, and D. A. Vignali. 2012. Loss of epigenetic modification driven by the Foxp3 transcription factor leads to regulatory T cell insufficiency. Immunity 36: 717–730.

67. Akimova, T., T. Zhang, D. Negorev, S. Singhal, J. Stadanlick, A. Rao, M. Annunziata, M. H. Levine, U. H. Beier, J. M. Diamond, J. D. Christie, S. M. Albelda, E. B. Eruslanov, and W. W. Hancock. 2017. Human lung tumor FOXP3+ Tregs upregulate four “Treg-locking” transcription factors. JCI Insight 2.

68. John, L. B., and A. C. Ward. 2011. The Ikaros gene family: transcriptional regulators of hematopoiesis and immunity. Mol Immunol 48: 1272–1278.

69. Kim, J., S. Sif, B. Jones, A. Jackson, J. Koipally, E. Heller, S. Winandy, A. Viel, A. Sawyer, T. Ikeda, R. Kingston, and K. Georgopoulos. 1999. Ikaros DNA-binding proteins direct formation of chromatin remodeling complexes in lymphocytes. Immunity 10: 345–355.

70. Josefowicz, S. Z., L. F. Lu, and A. Y. Rudensky. 2012. Regulatory T cells: mechanisms of differentiation and function. Annu Rev Immunol 30: 531–564.

71. Owen, D. L., L. E. Sjaastad, and M. A. Farrar. 2019. Regulatory T Cell Development in the Thymus. J Immunol 203: 2031–2041.

72. Raffin, C., L. T. Vo, and J. A. Bluestone. 2020. Treg cell-based therapies: challenges and perspectives. Nat Rev Immunol 20: 158–172.

73. Sharabi, A., M. G. Tsokos, Y. Ding, T. R. Malek, D. Klatzmann, and G. C. Tsokos. 2018. Regulatory T cells in the treatment of disease. Nat Rev Drug Discov 17: 823–844.

74. Liao, W., J. X. Lin, L. Wang, P. Li, and W. J. Leonard. 2011. Modulation of cytokine receptors by IL-2 broadly regulates differentiation into helper T cell lineages. Nat Immunol 12: 551–559.

75. Liao, W., R. Spolski, P. Li, N. Du, E. E. West, M. Ren, S. Mitra, and W. J. Leonard. 2014. Opposing actions of IL-2 and IL-21 on Th9 differentiation correlate with their differential regulation of BCL6 expression. Proc Natl Acad Sci U S A 111: 3508–3513.

76. Olson, M. R., F. F. Verdan, M. M. Hufford, A. L. Dent, and M. H. Kaplan. 2016. STAT3 Impairs STAT5 Activation in the Development of IL-9-Secreting T Cells. J Immunol 196: 3297–3304.

77. Liao, W., J. X. Lin, and W. J. Leonard. 2011. IL-2 family cytokines: new insights into the complex roles of IL-2 as a broad regulator of T helper cell differentiation. Curr Opin Immunol 23: 598–604.

78. Read, K. A., M. D. Powell, P. W. McDonald, and K. J. Oestreich. 2016. IL-2, IL-7, and IL-15: Multistage regulators of CD4(+) T helper cell differentiation. Exp Hematol 44: 799–808.

79. Leonard, W. J., J. X. Lin, and J. J. O’Shea. 2019. The gammac Family of Cytokines: Basic Biology to Therapeutic Ramifications. Immunity 50: 832–850.

