## Supplemental Figures for "Eos promotes T_H_2 differentiation by propagating the IL-2/STAT5 signaling pathway"

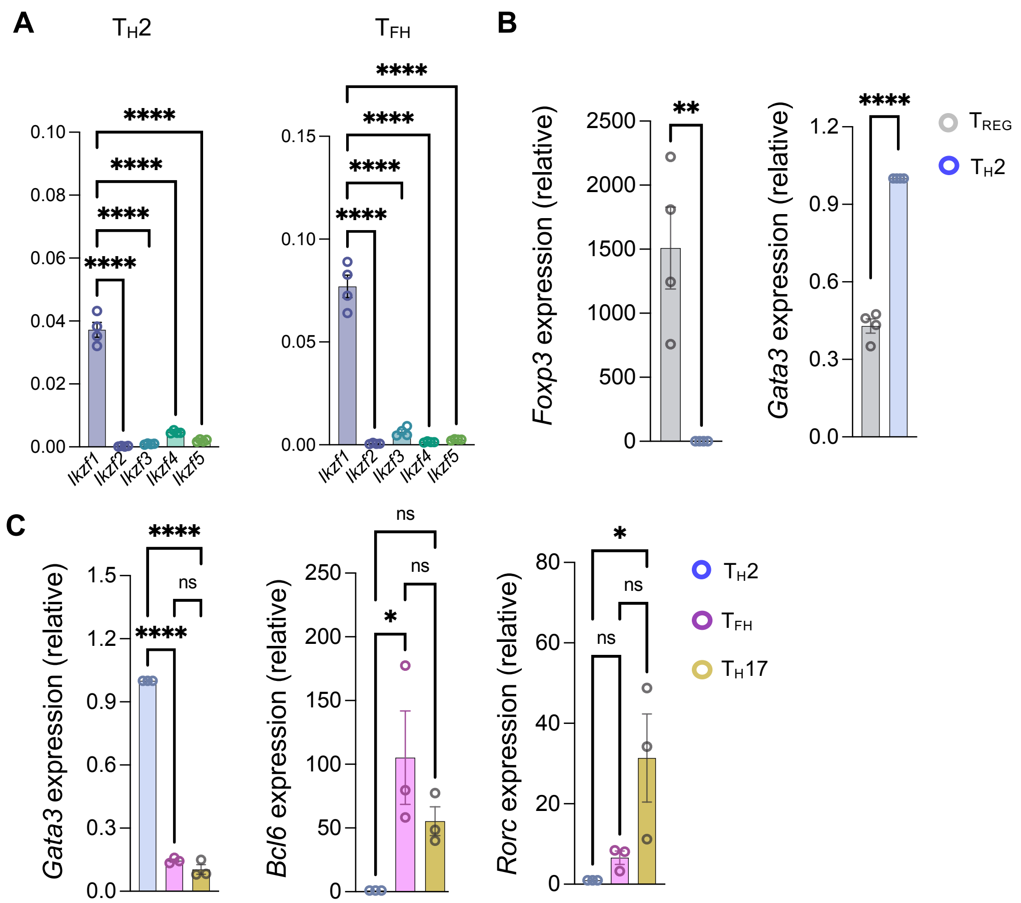


**Supplemental Figure 1**

**(A)** Naïve WT $CD4^+$ T cells from C57BL/6J mice were cultured with α-CD3/α-CD28 under $T_H2$- and $T_{FH}$-polarizing conditions for 3 days before being removed from stimulation and cultured in resting $T_H2$ (IL-4 and IL-2) and $T_{FH}$ (IL-6, IL-2) conditions for an additional 48 hours before harvesting on day 5. RNA was isolated and qRT-PCR was used to assess gene expression. Data were normalized to *Rps18* (n = 4 independent experiments, mean ± s.e.m., **** $P <$ 0.0001). **(B,C)** Cells were cultured as in 'A' under $T_{REG}$-, $T_H2$-, $T_{FH}$-, and $T_H17$-polarizing conditions. RNA was isolated and qRT-PCR was used to assess gene expression. Data were normalized to *Rps18* (n = 4 independent experiments, mean ± s.e.m., * $P < 0.05$, ** $P < 0.01$, *** $P < 0.001$, **** $P < 0.0001$; one-way ANOVA with Tukey's post-hoc test).

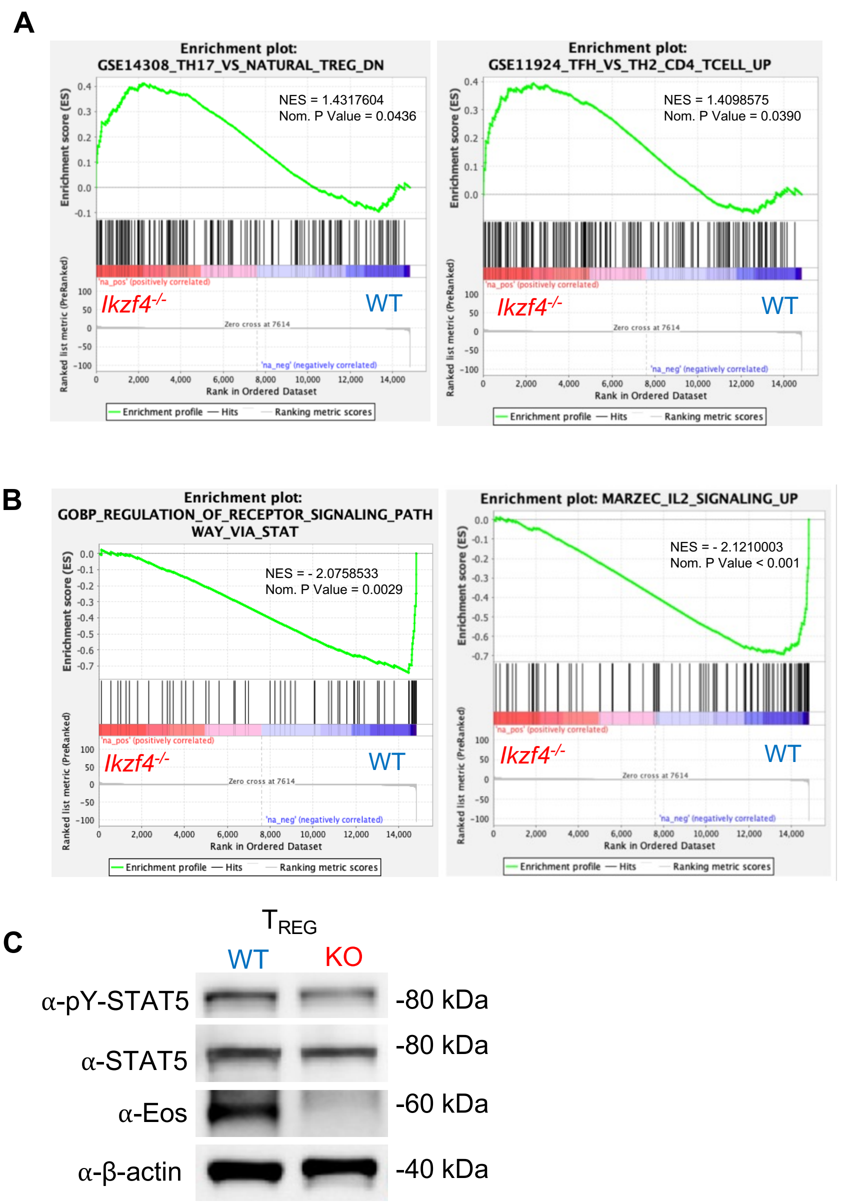


**Supplemental Figure 2**

**(A)** Naïve CD4$^+$ T cells from WT and Eos-deficient C57BL/6J mice were cultured on plates coated with α-CD3/α-CD28 stimulation under $T_H2$-polarizing conditions (**Table 1**) and harvested on day 3. RNA was isolated and qRT-PCR was used to assess gene expression. Data were normalized to *Rps18* and presented as fold change relative to the day 3 WT $T_H2$ sample (n = 7 independent experiments, mean ± s.e.m., * $P < 0.05$, ** $P < 0.01$, **** $P < 0.0001$; unpaired Student's *t*-test). **(B)** Pre-ranked (sign of fold change x -log$_{10}$(p-value)) genes were analyzed using the Broad Institute GSEA software for comparison against ‘immunological signature’, ‘gene ontology’, and ‘curated’ gene sets. Data are compiled from 3 biological replicates from 3 independent experiments. **(C)** Immunoblot analysis of pY-STAT5, STAT5B, and Eos protein expression in in vitro-generated $T_{REG}$ cells (n = 2 independent experiments).

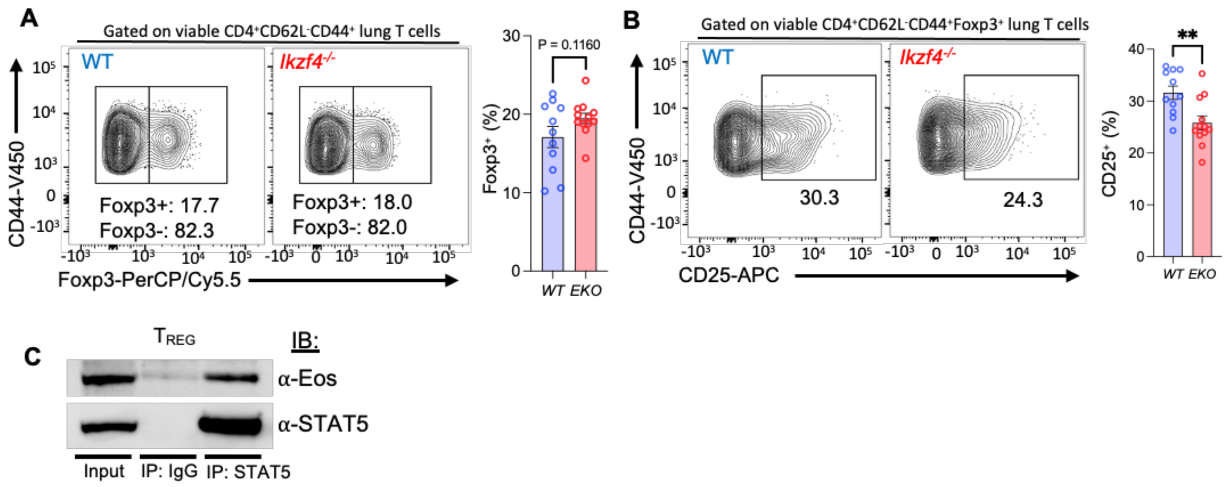


**Supplemental Figure 3**

**(A,B)** Single-cell suspensions from the lung were incubated in culture medium in the presence of PMA and ionomycin stimulation and protein transport inhibitors for 4 hours. Foxp3 protein expression (A) and CD25 production in CD4$^+$CD44$^+$CD62L$^-$Foxp3$^+$ populations (B) were analyzed via flow cytometry. Data are compiled from 3 independent experiments (n = 12 ± s.e.m; ***P* < 0.01; *****P* < 0.0001; unpaired Student's *t*-test). **(C)** Co-IP of in vitro $T_{REG}$ with immunoprecipitated STAT5 and probed for Eos on immunoblot (n = 3 independent experiments).
